# Maternal immune activation during pregnancy alters early neurobehavioral development in nonhuman primate offspring

**DOI:** 10.1101/2020.07.02.185363

**Authors:** Roza M. Vlasova, Ana-Maria Iosif, Amy M. Ryan, Takeshi Murai, Tyler A. Lesh, Douglas J. Rowland, Jeffrey Bennett, Casey E. Hogrefe, Richard J. Maddock, Michael J. Gandal, Daniel H. Geschwind, Cynthia M. Schumann, Judy Van de Water, A. Kimberley McAllister, Cameron S. Carter, Martin A. Styner, David G. Amaral, Melissa D. Bauman

## Abstract

**Background:** Human epidemiologic studies have implicated exposure to infectious or inflammatory insults during gestation in the etiology of neurodevelopmental disorders. Rodent models of maternal immune activation (MIA) have identified the maternal immune response as the critical link between maternal infection and aberrant brain and behavior development in offspring. The nonhuman primate MIA model provides an opportunity to maximize the translational utility of this model in a species more closely related to humans.

**Methods:** Here we evaluate the effects of MIA on brain and behavioral development in the rhesus monkey (*Macaca mulatta*). A modified form of the viral mimic, Polyinosinic-polycytidylic acid (PolyIC), was delivered to pregnant rhesus monkeys (*n*=14) in the late first trimester to stimulate a maternal immune response. Control dams received saline injections at the same gestational time points (*n*=10) or were untreated (*n*=4).

**Results:** MIA-treated dams exhibited a strong immune response as indexed by transient increases in sickness behavior, temperature and inflammatory cytokines. MIA-exposed offspring developed species typical milestones and demonstrate subtle changes in early in social development. However, magnetic resonance imaging demonstrated significant gray matter volume reductions in prefrontal and frontal cortices at 6, 12 and 24 months of age.

**Conclusions:** These findings provide new insights into the emergence of neuropathology in MIA-exposed primates and have implications for the pathophysiology of human psychiatric disorders associated with maternal gestational infection.

## INTRODUCTION

The need to understand the interaction between maternal exposure to infection during pregnancy and the subsequent increased risk of central nervous system (CNS) disorders in offspring has never been more urgent^1^. Children born to women exposed to infection have an increased risk of brain disorders with neurodevelopmental origins, including both schizophrenia (SZ) and autism spectrum disorder (ASD)^2,3^. The association between maternal infection and offspring neurodevelopmental disorders documented in epidemiological studies^4–7^ is further supported by the presence of infectious or inflammatory biomarkers in gestational biospecimens obtained from mothers of children later diagnosed with SZ or ASD^8–17^. The diversity of maternal infections associated with adverse offspring outcomes suggests that the maternal immune response may be the critical link between exposure to infection and altered fetal brain development. The maternal immune activation (MIA) hypothesis proposes that exposure to infection during pregnancy triggers an immune response that alters the maternal-fetal immune environment, disrupts fetal brain development, and serves as a “disease primer” for a range of neurodevelopmental and neuropsychiatric disorders in the offspring^18,19^.

Preclinical models provide a translational tool to systematically explore the etiological role of prenatal immune challenge in neurodevelopmental disorders^20^. MIA models use viral mimics, such as Polyinosinic:polycytidylic acid (Poly IC), to elicit an immune response during gestation and then characterize changes in the brain and behavioral development of the offspring in a controlled environment^21^. Rodents born to MIA-treated dams exhibit a number of alterations in brain and behavioral development that resemble features of human neurodevelopmental and neuropsychiatric disease^22^. For example, impairments in offspring sensory processing and social development are highly reproducible in the rodent MIA models in spite of methodological variability^23^. Moreover, cross-species MIA model comparisons in mice, rats and nonhuman primates provide new opportunities to maximize the translational utility of this promising animal model and to explore neurobiological mechanisms underlying human neurodevelopmental disorders^24–26^.

Our research team developed the first Poly IC-based nonhuman primate model to bridge the gap between rodent MIA models and human patient populations. In a previous study, pregnant rhesus monkeys (*Macaca mulatta*) injected with a modified form of Poly IC over four days at the end of either the first or second trimester exhibited a transient but potent immune response and produced offspring that developed abnormal repetitive behaviors, reduced affiliative vocalizations and altered immune regulation within the first two years^27,28^. As they matured, monkeys exposed to MIA in the first trimester also exhibited inappropriate interactions with novel conspecifics and failed to attend to salient social cues^29^. The presence of abnormal repetitive behaviors paired with alterations in social development exhibited by MIA-treated offspring suggests that circuity relevant to neurodevelopmental disorders, including both ASD and SZ, may be vulnerable to prenatal immune challenge. Although we are at the earliest stages of exploring the underlying neuropathology in the nonhuman primate MIA model, our preliminary neuroimaging studies using positron emission tomography revealed increased striatal dopamine in late adolescence^30^ and histopathological analysis revealed aberrant dendritic morphology in the dorsolateral prefrontal cortex (DLPFC)^31^. In the present study, we report developmental behavioral and neuroanatomical findings from a new and larger cohort of MIA-treated male offspring using longitudinal, multimodal neuroimaging paired with comprehensive behavioral phenotyping. We present the initial characterization of growth and development in this new cohort along with structural magnetic resonance imaging (MRI) data obtained at time roughly spanning early to mid-childhood in the rhesus monkey.

## METHODS

All experimental procedures were developed in collaboration with the veterinary, animal husbandry, and environmental enrichment staff at the California National Primate Research Center (CNPRC) and approved by the University of California, Davis Institutional Animal Care and Use Committee. All attempts were made (in terms of social housing, enriched diet, use of positive reinforcement strategies, and minimizing the duration of daily training/testing sessions) to promote normal social development and the psychological well-being of the animals that participated in this research. Detailed methods are provided in supplemental material.

### Animal selection

Pregnant dams were selected from the indoor, time-mated breeding colony based on age, weight, parity, and number of prior live births (Supplemental Table 1). Candidate dams between 5 and 12 years old carrying a male fetus were assigned to MIA (*n*=14) and control/saline (*n* =10) (Supplemental Table 2). Due to limited availability of male fetuses, untreated pregnant females confirmed to be carrying male fetuses were added to the control group (*n*=4). One offspring from the MIA group was euthanized at 6 months of age due to an unrelated health condition.

### Maternal immune activation and validation

Synthetic double-stranded RNA (polyinosinic:polycytidylic acid [Poly IC] stabilized with poly-L-lysine [Poly ICLC]) (Oncovir, Inc.; 0.25 mg/kg i.v.) or sterile saline (equivalent volume to Poly ICLC) was injected at 0730 hours in the cephalic vein in awake animals on gestational day (GD) 43, 44 and 46 (Supplemental Table 3). Health and behavioral observations were conducted three times pre-treatment, 6 hours after each of the three injections, and three times post-treatment. Blood was collected from the dams on approximately GD 40 while sedated for ultrasound, from awake animals on GD 44 and 46, six hours after Poly ICLC infusion, and on GD 51 or 52 while sedated for a recheck ultrasound for cytokine analysis. Blood samples were centrifuged and the serum was removed, aliquoted into 200uL samples, and frozen at −80°C until analysis. A longitudinal analysis on the maternal IL-6 response to Poly ICLC exposure was measured in serum collected at baseline (pre-exposure at GD 40) and after the second (GD 44) and third (GD 46) Poly ICLC injections using a nonhuman primate multiplexing bead immunoassay (Milipore-Sigma, Burlington, MA) that was analyzed using the flow-based Luminex™ 100 suspension array system (Bio-Plex 200; Bio-Rad Laboratories, Inc.).

### Rearing conditions and husbandry

Infants were raised in individual cages with their mothers, where they had visual access to other mother-infant pairs at all times. For 3 hours each day, one familiar adult male and four familiar mother-infant pairs were allowed to freely interact in a large cage (3m l x 1.8m w x 2m h) to facilitate species-typical social development. The infants were weaned from their mothers at 6 months of age and were permanently paired with a peer from their rearing group and in the same experimental group. Weanlings continued the same socialization routine through approximately 18 months of age.

### Offspring physical growth, neurodevelopmental milestones and early behavioral development

Measures of body growth (weight, crown rump length and head circumference) were collected at 1 and 3 months of age as well as at the neuroimaging time points (6, 12 and 24 months of age). The nonhuman primate offspring described in this paper are undergoing comprehensive assessments of social and cognitive development that will be the focus of future publications. Here we present data on a neonatal neurobehavioral assessment (1 week) and home cage observations of the mother-infant dyad (0-6 months), home cage observations with their age/sex/treatment matched cage mate (6-18 months) and introduction to a novel conspecific (11 months) (Supplemental Table 4). Behavioral observations were carried out by trained observers demonstrating an inter-observer reliability > 85% (agreements/ [agreements + disagreements] X 100). Each infant received a dye mark, allowing the observers to record behaviors while remaining blind to their experimental condition. Detailed methods are provided in our previous publications^27,32–34^ and in the supplemental material.

### Neuroimaging

Magnetic resonance imaging was performed under anesthesia at approximately 6, 12 and 24 months of age using a Siemens Magnetom Skyra 3-T (Davis, California) with 8-channel coil optimized for monkey brain scanning (RapidMR, Columbus, Ohio). T1 weighted images (480 sagittal slices) were acquired with TR=2500 ms, TE=3.65 ms, flip angle=7°, field of view 256×256, voxel size during acquisition 0.6×0.6×0.6 mm. Acquired images were interpolated during image reconstruction to 512×512 voxels with a final resolution of 0.3 × 0.3 × 0.3 mm. All images were analyzed by investigators who were blinded to group assignment. T1 weighted images were aligned into common atlas space^35^, bias field corrected, and brain masked using AutoSeg_3.3.2^36^. Brain masks were manually corrected if necessary. Following this preprocessing, T1 weighted images were segmented into gray matter (GM), white matter (WM) and cerebrospinal fluid (CSF) using NeosegPipeline_v1.0.8 ^37^(Supplemental Figure S1). Probabilistic tissue maps from structural multi-atlas templates were applied to each subject’s T1 weighted images via deformable registration. University of North Carolina (UNC) lobar parcellations were employed to parcellate the tissue segmentations into 24 lobar brain regions using the multi-atlas fusion in AutoSeg_3.3.2 (Supplemental Figure S1)^36^. For these regions, total GM and WM volumes were extracted. Lateral ventricle volumes were determined via semi-automated segmentation using the region competition deformable surface approach in ITK snap ^38^ as applied to the probability CSF maps from the tissue segmentation ^39^. In order to limit the number of comparisons, we selected five regions of interest based on a review of literature documenting brain alterations associated with MIA: prefrontal, frontal, cingulate, temporal limbic (including amygdala and hippocampus) regions (Supplemental Figure S2), and the lateral ventricles^40–44^.

### Statistical analysis overview

Statistical analysis was conducted using a general linear mixed models framework that can accommodate traditional general linear models (e.g., ANOVA and multiple linear regression) for data that were assumed independent across individuals, as well as mixed-effects linear models^45^ for data that was collected repeatedly for an individual (across time or conditions). An advantage of this approach is the ability to use all available data for an individual, to account for the effect of covariates of interest, and to directly model heterogeneous variances (across groups or conditions). Transformations were employed if assumptions of the linear models were not met and nonparametric techniques (exact Wilcoxon two-sample test) were used to compare groups when transformations were unsuccessful. All models were validated both graphically and analytically. Tests were two-sided, with *α* = 0.05. All analyses were conducted in SAS version 9.4. (SAS Institute Inc., Cary, NC). Detailed description of the specific statistical models used to analyze each of the offspring development, physical growth, behavioral, and neuroimaging data is available in Supplemental material.

## RESULTS

### Validation of maternal immune activation

Blood samples collected six hours after the second (GD 44) and third (GD 46) Poly ICLC injections confirmed a strong pro-inflammatory cytokine response as indexed by change in IL-6 from baseline samples (Table 1; Figure S4). Dams that received Poly ICLC injections also exhibited sickness behaviors, including reduced appetite and fever (Supplemental Tables 5-6 and Figure S3).

**Table 1.** Maternal IL-6 response.

|  | MIA<br>(*n* = 14) |  | Control<br>(*n* = 10) |  |
| --- | --- | --- | --- | --- |
|  | Mean (*SD*) | Median [Range] | Mean (*SD*) | Median [Range] |
| ***IL-6 (pg/ml)*** |  |  |  |  |
| Pre-dosing (GD 40) | 10.9 (28.7) | 0.9 [0.0 – 108.3] | 0.6 (0.6) | 0.4 [0.0 – 1.8] |
| After second injection (GD 44) | 777.9 (1068.6) | 450.9 [90.8 – 4341.7] | 4.6 (9.6) | 1.0 [0.0 – 31.4] |
| After third injection (GD 46) | 493.0 (751.4) | 130.4 [51.7 – 2780.7] | 6.8 (18.6) | 0.4 [0.0 – 59.5] |
| ***IL-6 change from baseline (pg/ml)*** |  |  |  |  |
| After second injection (GD 44) | 767.3 (1072.9) | 450.9 [89.1 – 4341.7] | 4.0 (9.4) | 0.6 [-0.7 – 30.4] |
| After third injection (GD 46) | 482.3 (754.0) | 129.8 [50.0 – 2777.7] | 6.2 (18.6) | 0.0 [-0.2 – 59.1] |

Abbreviations: MIA, maternal immune activation; SD, standard deviation; GD, gestation day.

### Offspring development

There were no significant differences between MIA-treated and control offspring in growth trajectories (Figure 1a, b, c, d, Supplemental Table 7), neuromotor-reflexes (Supplemental Table 8) or home cage check list observations of mother-infant and familiar infant-infant interactions (Supplemental Tables 9, 10). Subtle group differences were detected when we evaluated offspring response to a novel age/sex matched conspecific at 11 months of age (Figure 2). Table 2 summarizes the frequency and duration of the behaviors for the two groups and the results of the general linear models used to analyze them after accounting for age at testing (centered at median, 10.2 months) and stimulus animal. For most of the behaviors, there was evidence of a group by age at testing interaction. MIA infants exhibited similar durations in nonsocial and social chamber regardless of age, while older control animals tended to have longer duration in nonsocial chamber and shorter durations in the social chamber than younger animals. This pattern resulted in subtle shorter duration in nonsocial chamber and longer duration in social chamber in the MIA-treated animals than in same age controls at 10.2 months (*p* = .07 and .02, respectively). The same pattern of subtle group differences at older ages was present for duration of contact and proximity with stimulus animal and frequency of proximity (*p* = .07, .03, and .08, respectively for group differences at 10.2 months). MIA animals had consistently more entries in the social chamber (*p* = .09), regardless of age.

**Figure 1.**
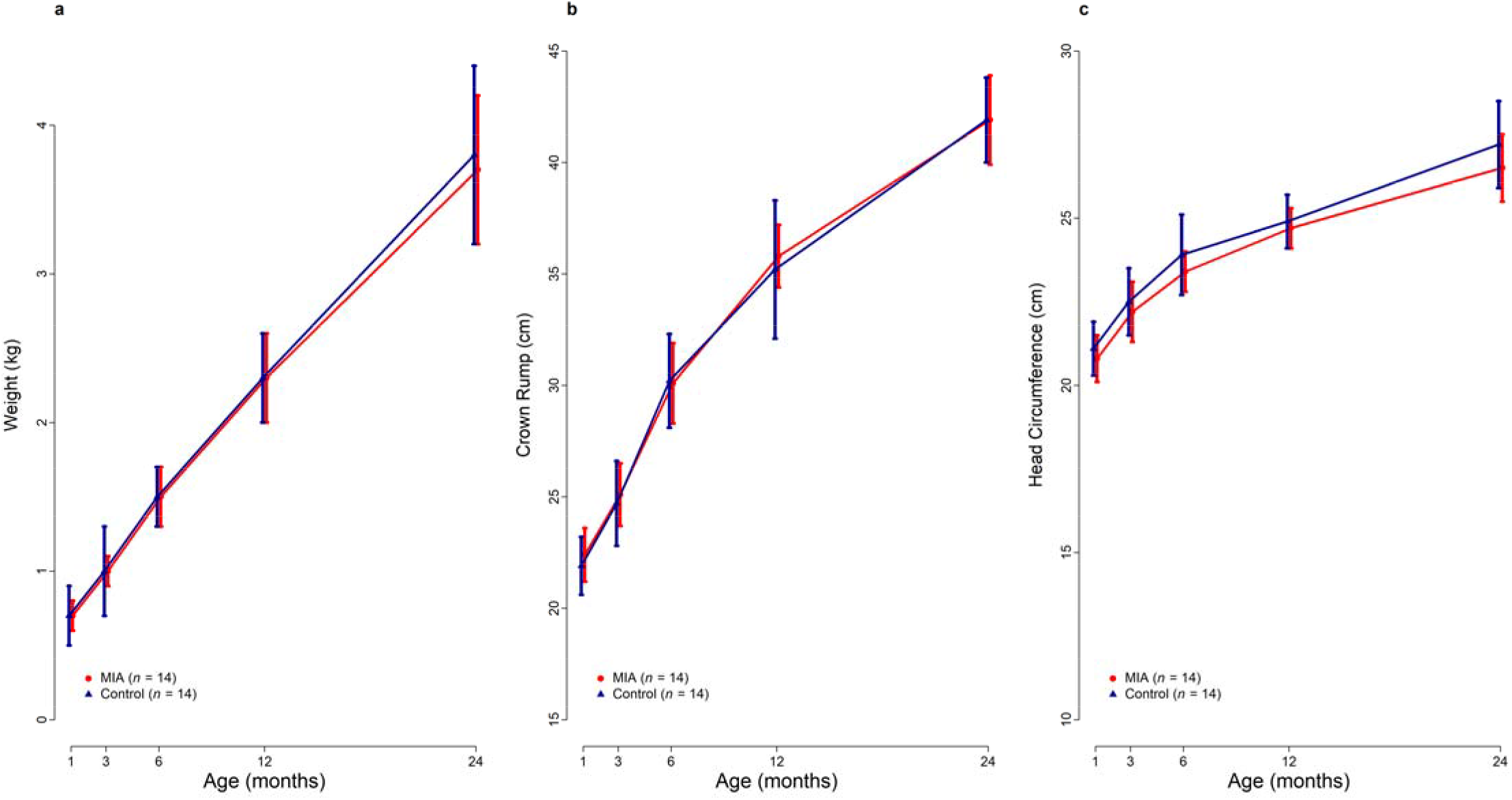
Average trajectory for weight (a), crown rump length (b) and head circumference (c) for the MIA and Control animals from 1 month through 24 months. Vertical bars represent 1 standard deviation.

**Figure 2.**
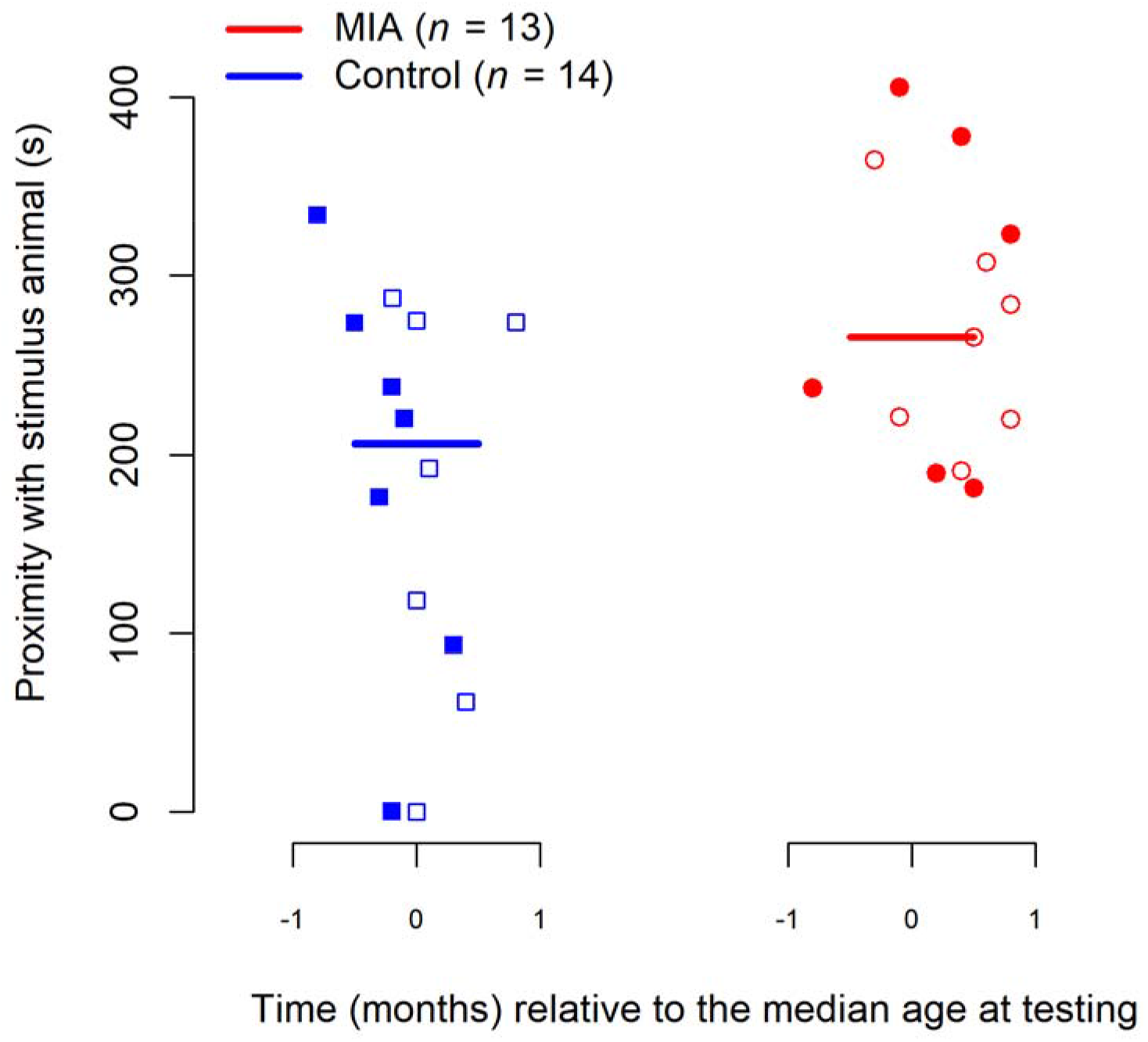
Duration for proximity with stimulus animal by group. Animals were tested between 9.4 and 11 months (median =10.2) using two same-sex stimulus animals (represented by filled or hollow symbols). Horizontal lines represent group medians.

**Table 2.**
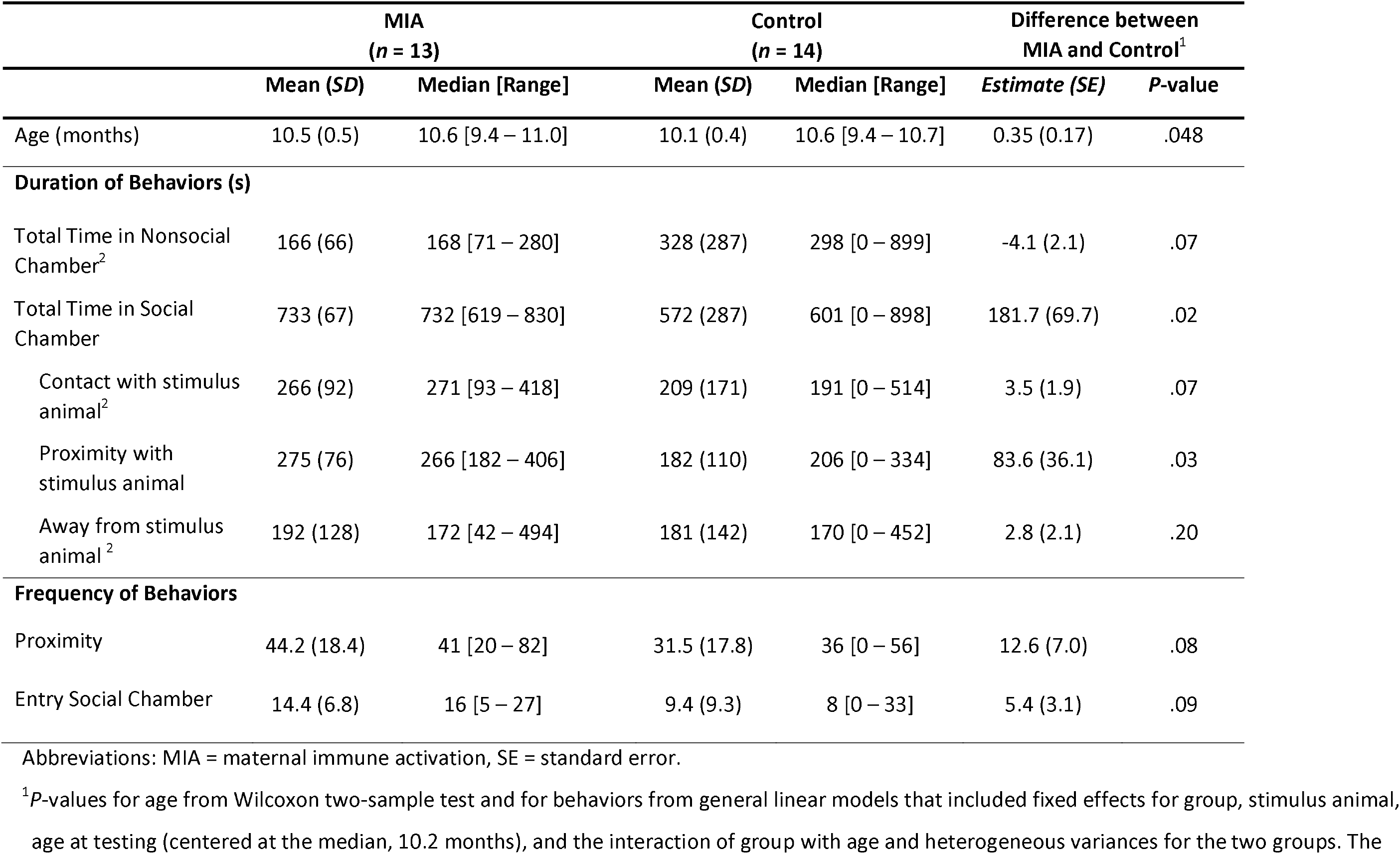
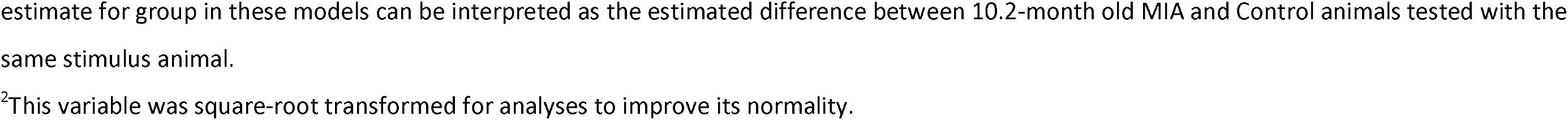
Social Approach Paradigm. Summary of duration and frequency of behaviors for the MIA and Control animals and parameter estimates and standard errors of the general linear models assessing the relationship between group and duration and frequency of behavior variables

|  | MIA<br>(*n* = 13) |  | Control<br>(*n* = 14) |  | Difference between<br>MIA and Control<sup>1</sup> |  |
| --- | --- | --- | --- | --- | --- | --- |
|  | Mean (*SD*) | Median [Range] | Mean (*SD*) | Median [Range] | Estimate (*SE*) | *P*-value |
| Age (months) | 10.5 (0.5) | 10.6 [9.4 – 11.0] | 10.1 (0.4) | 10.6 [9.4 – 10.7] | 0.35 (0.17) | .048 |
| **Duration of Behaviors (s)** |  |  |  |  |  |  |
| Total Time in Nonsocial Chamber<sup>2</sup> | 166 (66) | 168 [71 – 280] | 328 (287) | 298 [0 – 899] | -4.1 (2.1) | .07 |
| Total Time in Social Chamber | 733 (67) | 732 [619 – 830] | 572 (287) | 601 [0 – 898] | 181.7 (69.7) | .02 |
| Contact with stimulus animal<sup>2</sup> | 266 (92) | 271 [93 – 418] | 209 (171) | 191 [0 – 514] | 3.5 (1.9) | .07 |
| Proximity with stimulus animal | 275 (76) | 266 [182 – 406] | 182 (110) | 206 [0 – 334] | 83.6 (36.1) | .03 |
| Away from stimulus animal<sup>2</sup> | 192 (128) | 172 [42 – 494] | 181 (142) | 170 [0 – 452] | 2.8 (2.1) | .20 |
| **Frequency of Behaviors** |  |  |  |  |  |  |
| Proximity | 44.2 (18.4) | 41 [20 – 82] | 31.5 (17.8) | 36 [0 – 56] | 12.6 (7.0) | .08 |
| Entry Social Chamber | 14.4 (6.8) | 16 [5 – 27] | 9.4 (9.3) | 8 [0 – 33] | 5.4 (3.1) | .09 |

Abbreviations: MIA = maternal immune activation, SE = standard error.
<sup>1</sup>*P*-values for age from Wilcoxon two-sample test and for behaviors from general linear models that included fixed effects for group, stimulus animal, age at testing (centered at the median, 10.2 months), and the interaction of group with age and heterogeneous variances for the two groups. The
estimate for group in these models can be interpreted as the estimated difference between 10.2-month old MIA and Control animals tested with the same stimulus animal.
<sup>2</sup>This variable was square-root transformed for analyses to improve its normality.

### Neuroimaging

Table 3 summarizes the volumetric measures for the two groups and Supplemental Table 11 displays the results of the mixed-effects linear models for global measures. The two groups had parallel growth trajectories from 6 to 24 months on all global measures i.e. total brain volume, total gray matter volume and total white matter volume; none of the group by time interactions reached statistical significance. Total brain volume from 6 to 24 months for the MIA-treated animals was consistently smaller than for the control animals although the difference was not significant (estimated difference [est.] = −5504 mm^3^, *p* = .12). The same pattern of lower levels in the MIA-treated animals than in the controls was present in total gray matter (est. = −3862 mm^3^, *p* = .11).

**Table 3.** Summary for the GM and WM volumetric measures (mm^3^) from 6 to 24 months.

|  | 6 Months |  | 12 Months |  | 24 Months |  |
| --- | --- | --- | --- | --- | --- | --- |
|  | MIA | Control | MIA | Control | MIA | Control |
|  | (n = 13) | (n = 14) | (n = 13) | (n = 14) | (n = 13) | (n = 14) |
| *Age (days) at scan, mean (SD) [Range]* |  |  |  |  |  |  |
|  | 180 (2) | 181 (5) | 365 (2) | 365 (1) | 730 (1) | 729 (1) |
|  | [177-182] | [177-199] | [364-369] | [364-366] | [729-732] | [729-733] |
| *Global Measures, Mean (SD)* |  |  |  |  |  |  |
| TBV | 81429 (8113) | 86200 (7724) | 83522 (9692) | 88781 (8550) | 87821 (10381) | 94268 (9060) |
| GM | 59665 (6002) | 63271 (5146) | 59404 (6884) | 63073 (5538) | 60710 (7240) | 65058 (5710) |
| WM | 21764 (2245) | 22929 (2666) | 24118 (2889) | 25708 (3133) | 27111 (3270) | 29210 (3598) |
| Left V | 227 (51) | 263 (112) | 228 (64) | 264 (142) | 246 (74) | 312 (147) |
| Right V | 296 (179) | 258 (82) | 309 (230) | 245 (72) | 340 (313) | 285 (73) |
| *Frontal Measures, Mean (SD)* |  |  |  |  |  |  |
| Left GM | 3397 (347) | 3740 (253) | 3439 (431) | 3814 (335) | 3563 (463) | 4043 (365) |
| Right GM | 3362 (328) | 3693 (275) | 3414 (385) | 3780 (332) | 3582 (451) | 4025 (374) |
| Left WM | 1308 (141) | 1411 (167) | 1453 (184) | 1586 (191) | 1710 (190) | 1854 (219) |
| Right WM | 1279 (166) | 1364 (167) | 1446 (169) | 1566 (193) | 1649 (203) | 1828 (199) |
| *Prefrontal Measures, Mean (SD)* |  |  |  |  |  |  |
| Left GM | 3165 (449) | 3570 (339) | 3151 (484) | 3547 (379) | 3146 (501) | 3588 (389) |
| Right GM | 3148 (439) | 3557 (321) | 3126 (466) | 3531 (381) | 3153 (490) | 3581 (400) |
| Left WM | 779 (140) | 840 (119) | 918 (165) | 1003 (138) | 1070 (192) | 1196 (167) |
| Right WM | 788 (131) | 862 (123) | 913 (161) | 1014 (147) | 1081 (186) | 1212 (174) |
| *Cingulate Measures, Mean (SD)* |  |  |  |  |  |  |
| Left GM | 1061 (132) | 1138 (120) | 1052 (131) | 1152 (136) | 1053 (150) | 1145 (135) |
| Right GM | 1091 (140) | 1177 (123) | 1084 (144) | 1185 (119) | 1084 (154) | 1187 (131) |
| Left WM | 169 (24) | 191 (33) | 186 (25) | 209 (36) | 212 (32) | 242 (42) |
| Right WM | 162 (26) | 179 (27) | 175 (31) | 196 (32) | 204 (35) | 229 (38) |
| *Temporal Limbic Measures, Mean (SD)* |  |  |  |  |  |  |
| Left GM | 1174 (87) | 1193 (111) | 1278 (112) | 1327 (124) | 1429 (144) | 1460 (142) |
| Right GM | 1241 (113) | 1249 (109) | 1354 (133) | 1411 (125) | 1500 (178) | 1557 (131) |
| Left WM | 212 (21) | 222 (24) | 227 (30) | 237 (32) | 275 (32) | 278 (36) |
| Right WM | 216 (28) | 220 (27) | 236 (24) | 245 (27) | 258 (36) | 285 (35) |

Abbreviations: TBV, total brain volume; GM, gray matter; WM, white matter; V, Ventricle.

Supplemental Table 12 summarizes the results of the unadjusted analyses for gray and white matter volumes in the five regions of interest. For gray matter, the two groups had parallel developmental trajectories from 6 to 24 months in all four regions; none of the group-by-time interactions reached significance. Yet, for the gray matter in the frontal and prefrontal regions (Figure 3), MIA-treated monkeys had lower values than controls at 6 months and these differences persisted at later ages (Frontal: est. = −382.6 mm^3^, *p* = .009, Prefrontal: est. = −414.5 mm^3^, *p* = .02; Supplemental Table 12). Table 4 summarizes the results of the analyses after adjusting for total brain volume. The magnitude of the group differences decreased, but remained significant, in both frontal and prefrontal gray matter (Frontal: est. = −155.3 mm^3^, *p* = .02, Prefrontal: est. = −184.7 mm^3^, *p* = .02).

**Figure 3.**
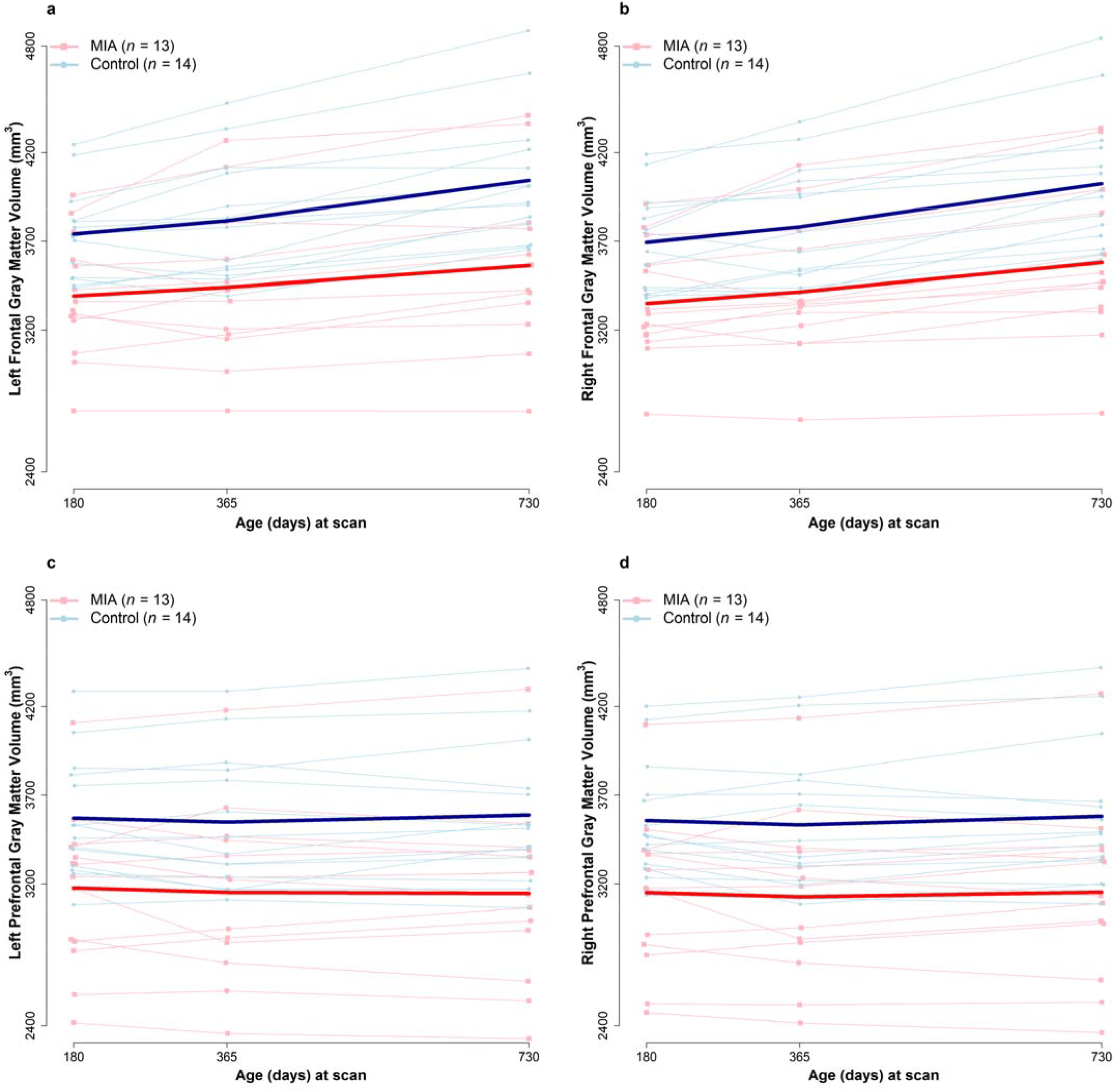
Gray matter volume trajectories in MIA and Control for left and right Frontal (panels a, b) and Prefrontal (panels c, d) regions. The light lines represent individual trajectories and dark lines represent average values for gray matter in the two groups.

**Table 4.** Parameter estimates from the linear mixed-effects models for gray and white ROI volumetric measures adjusted for total brain volume

| Gray Matter Volume |  |  |  |  |  |  |  |  |
| --- | --- | --- | --- | --- | --- | --- | --- | --- |
| Model term | Frontal |  | Prefrontal |  | Cingulate |  | Temporal Limbic |  |
|  | Estimate (SE) | P | Estimate (SE) | P | Estimate (SE) | P | Estimate (SE) | P |
| Intercept | 3709.8 (41.8) | <.001 | 3547.6 (50.5) | <.001 | 1176.5 (14.6) | <.001 | 1266.0 (15.1) | <.001 |
| Difference (mm<sup>3</sup>) *MIA vs. Control* | **-155.3 (61.0)** | **.02** | **-184.7 (73.7)** | **.02** | -23.9 (20.9) | .26 | 33.5 (20.6) | .12 |
| Difference (mm<sup>3</sup>) Time 2 vs. Time 1 | -31.9 (17.2) | .07 | -119.9 (15.1) | <.001 | -28.5 (5.8) | <.001 | 98.1 (9.3) | <.001 |
| Difference (mm<sup>3</sup>) Time 3 vs. Time 1 | -37.4 (26.7) | .17 | -298.1 (26.2) | <.001 | -91.7 (9.0) | <.001 | 178.9 (11.9) | <.001 |
| Difference (mm<sup>3</sup>) Left vs. Right | 24.0 (13.4) | .09 | 11.9 (7.4) | <.001 | -34.5 (9.2) | <.001 | -75.3 (7.3) | <.001 |
| Brain volume (cm<sup>3</sup>) | 41.8 (2.9) | <.001 | 41.8 (3.1) | <.001 | 12.7 (1.0) | <.001 | 12.7 (1.1) | <.001 |
| White Matter Volume |  |  |  |  |  |  |  |  |
| Model term | Frontal |  | Prefrontal |  | Cingulate |  | Temporal Limbic |  |
|  | Estimate (SE) | P | Estimate (SE) | P | Estimate (SE) | P | Estimate (SE) | P |
| Intercept | 1381.8 (20.3) | <.001 | 870.3 (14.5) | <.001 | 180.4 (3.4) | <.001 | 222.8 (4.2) | <.001 |
| Difference (mm<sup>3</sup>) *MIA vs. Control* | -29.0 (28.0) | .32 | -12.6 (21.0) | .56 | -5.2 (4.9) | .30 | 4.3 (5.9) | .47 |
| Difference (mm<sup>3</sup>) Time 2 vs. Time 1 | 130.8 (9.8) | <.001 | 99.8 (7.3) | <.001 | 8.6 (1.9) | <.001 | 12.5 (1.8) | <.001 |
| Difference (mm<sup>3</sup>) Time 3 vs. Time 1 | 290.9 (14.0) | <.001 | 203.2 (10.3) | <.001 | 24.2 (2.6) | <.001 | 36.5 (2.7) | <.001 |
| Difference (mm<sup>3</sup>) Left vs. Right | 31.8 (7.9) | <.001 | -10.7 (3.9) | .01 | 10.4 (2.2) | <.001 | -1.6 (2.1) | .46 |
| Brain volume (cm<sup>3</sup>) | 17.8 (1.5) | <.001 | 16.4 (1.1) | <.001 | 3.1 (0.3) | <.001 | 2.7 (0.3) | <.001 |

Note: MIA = maternal immune activation, SE = standard error, Time 1 = 6 months, Time 2 = 12 months, Time 3 = 24 months. Mixed-effects linear regression models were fitted to 13 MIA and 14 control animals and included fixed effects for group, time and their interaction, side, brain volume, and a random effect for animal. Interactions were not retained in the reported models since the overall tests for time by group were not significant. Brain volume was centered at 86,200; thus, the intercept can be interpreted as the predicted Time 1 right volume (in mm<sup>3</sup>) for a *Control* animal with a brain of 86,200 mm<sup>3</sup>. The estimate for brain volume can be interpreted as the average increase in ROI volume (in mm<sup>3</sup>) for a 1 cm<sup>3</sup> increase in brain volume. Statistically significant MIA vs. Control group differences ( $p < .05$ ) bolded for emphasis.

For both frontal and prefrontal white matter, there was an interaction between time and group in unadjusted analyses. The two groups had similar levels of white matter at 6 months, but the control group tended to have larger volume increases over time, resulting in higher volumes at 24 months than the MIA group (Frontal: est. = −163.5 mm^3^, *p* = .03, Prefrontal: est. = −128.1 mm^3^, *p* = .04). These differences did not persist after adjusting for total brain volume.

## DISCUSSION

The MIA model has emerged as a translational tool to explore the developmental pathophysiological mechanisms that link exposure to maternal infection during pregnancy with an increased risk of neurodevelopmental disorders in the offspring^20^. Although the vast majority of MIA model research has been carried out in rodents, the nonhuman primate provides a unique opportunity to evaluate the impact of MIA on brain and behavioral development in a species more closely related to humans. Our previous nonhuman primate MIA model work demonstrated that rhesus monkeys exposed to prenatal immune challenge exhibit alterations in brain and behavioral development relevant to human neurodevelopmental disorders^27–29^. Here we present initial results from a new cohort of MIA-treated nonhuman primate offspring undergoing longitudinal neuroimaging as part of a comprehensive assessment of brain and behavioral development. Structural MRI data, obtained at 6, 12 and 24 months of age, revealed deviations from species-typical brain development trajectories in the MIA-treated offspring. Compared with controls, the MIA-treated offspring exhibited reductions in frontal grey matter volumes at each time point, which roughly spans the period of early to middle human childhood. During this time, the MIA-treated offspring did not differ from controls on early assessments of physical growth and development, neonatal reflexes or preliminary evaluations of interactions with the mothers or familiar rearing partners. However, the MIA-treated offspring did exhibit subtle alterations in species-typical social behavior when introduced to a novel conspecific at approximately 11 months of age. These results provide evidence of early volumetric reductions paired with subtle alterations in social development following prenatal immune challenge in the rhesus monkey and extend the results from rodent MIA models to a primate species.

Gestational timing, choice of species, source of immune activating agent and subsequent magnitude of the maternal immune activation determine the impact on offspring neurodevelopment in preclinical MIA models^23^. Here we utilize a Poly IC based nonhuman primate MIA model to stimulate an inflammatory cytokine response late in the first trimester that mimics a moderate to severe infection during pregnancy. MIA-treated dams exhibited a transient immune response characterized by fever, reduced appetite and elevated inflammatory cytokines, including IL-6, which is necessary and sufficient for MIA to alter brain development and behavior in rodent offspring^46^. To our knowledge, this is the only nonhuman primate MIA model using Poly ICLC to stimulate the maternal immune response early in gestation, though other nonhuman primate prenatal immune challenge models have reported alterations in offspring brain and behavioral development following exposure to influenza^43^ or bacterial endotoxins in the third trimester^44^. Interestingly, similar to the MIA-treated offspring in the present study, the influenza exposed offspring exhibited early reductions in grey matter volume in the absence of overt behavioral changes^43^.

Volumetric reductions have also been noted in rodent MIA models^47–51^. Weiner and colleagues first documented longitudinal region-, age- and sex-specific changes in brain development in MIA-treated rat offspring using manual segmentation of regions of interest to quantify volumetric changes from postnatal day (PND) 35-90^52^. MIA-treated offspring had smaller absolute volumes of the hippocampus, striatum and prefrontal cortex and larger lateral ventricles compared to controls, though sex-specific trajectories were observed. Vernon and colleagues recently reported age-specific volumetric reductions in the anterior cingulate cortex and hippocampus of MIA-offspring using a semi-automated technique and volumetric reductions in additional cortical areas, including prefrontal cortex, using a fully automated technique to explore longitudinal changes in MIA-treated rat offspring^40^. Collectively, the rodent neuroimaging studies suggest that MIA is associated with volumetric reductions, though these changes are age, region, and sex-specific^22^. Although comparing developmental trajectories across species is challenging, it is noteworthy that reduced frontal volume has been detected in both mid-gestation MIA-treated rats^22,40^ and late first trimester MIA-treated nonhuman primates.

Rodent MIA models have identified numerous changes in neuronal migration, number and density, as well as alterations in dendritic structure and synapse formation that could contribute to aberrant brain growth trajectories (for review^53^). Although we are at the earliest stages of exploring neuropathology in the nonhuman primate MIA model, the gestational timing of the prenatal immune challenge may provide some insight into which stages of fetal brain development may be affected. In the present study, dams received Poly ICLC injections on GD 43, 44, and 46, which corresponds to late first trimester (GD 0-55) of the 165 day macaque gestation. Peak periods of neurogenesis for subcortical structures, including the amygdala^54^ and thalamus^55^, as well as the early stages of neurogenesis for the striatum^56^ and hippocampus^57^ occur during this time. In macaques, the early stages of corticogenesis also begin at the end of the first trimester and continue through the second trimester (GD 56-110)^58^. Exposure to prenatal immune challenge on GD 43-46 is positioned to disrupt the finely orchestrated events of neurodevelopment taking place near the end of the first trimester and could initiate a cascade of aberrant brain and behavioral development. Our preliminary evaluation of brain tissue from our earlier cohort of MIA-treated monkeys revealed subtle alterations in dendritic morphology in the DLPFC during adolescence^31^. It is not known if these subtle morphological changes in the DLPFC are present early in development or emerge as the MIA-treated offspring mature. However, the reduction in frontal cortex volume reported in the present MRI study suggests that the frontal cortex of MIA-treated offspring deviates from typical brain development trajectories in the early postnatal period and may be a particularly vulnerable region.

An emerging view is that activation of the maternal immune system during pregnancy serves as a “disease primer” that, in combination with other genetic and/or environmental risk factors, may increase the risk for specific neurodevelopmental disorders^18^, including both ASD and SZ^24,59^. While comparisons between animal models and clinical disorders must be made with caution, the nonhuman primate model does more closely approximate humans in maternal immune response, placental function and offspring brain and behavioral development^25^. We have previously demonstrated that rhesus monkeys exposed to prenatal immune challenge develop aberrant behaviors after a period of early typical development^27–29^ and exhibit increased striatal dopamine^30^, which is a hallmark molecular biomarker of SZ^60^. The reductions in cortical grey matter volume observed in this new cohort of MIA-treated monkeys also align with neuroimaging studies from adolescents and young adults diagnosed with SZ^61–71^, though at a much earlier time in postnatal development. While it is incredibly challenging to carry out adequately powered prospective neuroimaging studies in infants at risk for SZ^72^, there is ample evidence from cohort studies of behavioral and cognitive changes in young children who go on to develop SZ^73–75^. Changes in early postnatal brain development have also been documented in other neurodevelopmental disorders associated with maternal infection, such as ASD. Early brain overgrowth has been consistently reported in young children with ASD^76–78^ but findings are less consistent in older individuals. Longitudinal evaluation spanning early childhood through adolescence and adulthood are needed to better understand alterations in brain growth trajectories of both ASD and SZ.

Across species, offspring born to MIA-treated dams exhibit alterations in brain and behavioral development relevant to human neurodevelopmental and neuropsychiatric disease and supported the initial interpretation of the MIA model as an animal model “of” ASD or SZ. As the use of preclinical models continues to evolve towards a hypothesis-based approach^79^, the MIA model has been highlighted in recently released NIH-guidelines (NOT-MH-19-053) as a model “for” examining the effects of prenatal immune challenge on offspring neurodevelopment. The MIA-treated monkeys in the current study did not differ from controls on measures of growth, neuromotor reflex development or interactions with their mothers of familiar rearing partners. However, subtle differences were detected in the MIA-treated animals when introduced to an unfamiliar animal at approximately 1 year of age using a modified version of the rodent 3-chamber social approach assay. The MIA-treated monkeys spent more time than controls in the large “social chamber” that contained an unfamiliar age/sex-matched conspecific temporarily confined to a small holding cage. While in the social chamber, the MIA monkeys spent more time than controls in immediate proximity (i.e., within arm’s reach) to the unfamiliar animal and demonstrated trend level differences in the amount of time in contact with the unfamiliar animal’s holding cage. For primates, the decision to approach and interact with another animal depends on a number of factors, including individual temperament differences paired with physical characteristics and behavior of the unfamiliar animal and the contact of the encounter^80–85^. For many species of nonhuman primates, approaching an unfamiliar conspecific or behaving impulsively with familiar animals can be met with negative outcomes such as physical aggression^86–92^. Interestingly, a similar pattern of behavior was observed in our previous pilot cohort of MIA-treated monkeys at 2 years of age^27^. The emergence of atypical social behavior in MIA-treated animals at approximately 1 year of age suggests that the alterations in species-typical social behavior that are highly reproducible in the rodent MIA model^23^ are also present in the nonhuman primate model. This will allow for the comparison of MIA downstream effects on evolutionarily conserved behavioral and biological outcome measures^93–95^.

The results of the present study extend the results of rodent MIA models into a species more closely related to humans. Limitations of the current nonhuman primate study include a relatively modest sample size and the exclusion of female offspring. We recognize that sex differences are emerging as a critical factor in MIA model studies^96^, and plan to next explore the nonhuman primate MIA model in female offspring. Although subtle group differences were detected at these early ages, a more comprehensive evaluation of social and cognitive development is underway along with continued longitudinal neuroimaging to characterize the emergence of neurobehavioral alterations in MIA-exposed primates. Our observation of reduced cortical volume in MIA-treated monkeys paired with subtle behavioral changes contributes to mounting evidence that early exposure to prenatal immune challenge triggers a cascade of brain and behavioral changes that are highly relevant to human neurodevelopmental and psychiatric disorders. Additional clinical and translational research is needed to understand how other factors, including genetic risk, gestational timing, nature and intensity of the maternal immune response and additional postnatal events determine which (if any) disease phenotype results from prenatal exposure to MIA.

## Acknowledgments

These studies were supported by the UC Davis Conte Center to CSC (NIMH; P50MH106438). Development of the nonhuman primate model and behavioral characterization of the offspring were supported by (P50MH106438-6618) to MDB. Neuroimaging studies were supported by (P50MH106438-6616) to DGA. Cytokine analysis was supported by Biological Analysis Core of the MIND Institute Intellectual and Developmental Disabilities Research Center (U54 HD079125). AR was supported by the UC Davis Autism Research Training Program (T32MH073124). Additional support provided by the base grant (RR00169) of the California National Primate Research Center (CNPRC). We thank the veterinary and animal services staff of the CNPRC for care of the animals. Poly ICLC was kindly provided by Dr. Andres Salazar, MD, Oncovir, Washington D.C.

