## Supplemental Material for "Maternal immune activation during pregnancy alters early neurobehavioral development in nonhuman primate offspring"

**SUPPLEMENTAL METHODS**

**Animal selection.**

Pregnant dams were selected from the indoor time-mated breeding colony at the California National Primate Research Center (CNPRC) based on previous social housing experience, positive indoor pairing history, experience rearing at least one infant to weaning, and were not in any prior immune-related studies (Supplemental Table S1). Fetal sex was determined through maternal blood samples ^1^ by approximately gestational day (GD) 40, and only dams carrying male fetuses were eligible for the study. Candidate dams carrying a male fetus between 5 and 12 years old and were assigned to MIA (*n*=14) and control/saline (*n*=10). Pregnancy viability was then confirmed by ultrasound at GD 40, GD 51-52, GD 100 and GD 150. Due to limited availability of male fetuses, untreated females were added to the control group (*n*=4). All females remained in their original housing locations and with their established pair-mate throughout pregnancy and were relocated to the single project housing room after offspring were born. One offspring from the MIA group was euthanized at 6 months of age due to an unrelated health condition.

**Supplemental Table S1.** Summary of dam characteristics.

|  | **MIA**  **(*n* = 14)** | **Control**  **(*n* = 10 saline, *n* = 4 untreated)** |
| --- | --- | --- |
| ***Dam Characteristic*** | **Mean (*SD*)** | **Mean (*SD*)** |
| Age at Conception (years) | 9.23 (2.36) | 8.69 (2.18) |
| Weight at GD 40 (kg) | 7.50 (1.45) | 7.74 (1.63) |
| Prior Conceptions | 4.71 (2.02) | 3.64 (2.24) |

Abbreviations: MIA, maternal immune activation; SD, standard deviation; GD, gestation day.

**Experimental Groups.**

Offspring included MIA (*n*=14) and control (*n*=10 saline treated and *n*=4 untreated) groups (Supplemental Table S2). One offspring from the MIA group was euthanized at 6 months of age due to an unrelated health condition and is not included in behavior or neuroimaging data sets after 6 months of age.

**Supplemental Table S2.** Experimental Groups

| **Experimental Group** | **Group Size** |
| --- | --- |
| MIA | *n*=14 males |
| Saline Controls | *n*=10 males |
| Untreated Controls | *n*=4 males |

**Maternal immune activation.**

Synthetic double-stranded RNA (polyinosinic:polycytidylic acid [Poly IC] stabilized with poly-L-lysine [Poly ICLC]) (Oncovir, Inc.; 0.25 mg/kg i.v.) or sterile saline (equivalent volume to Poly ICLC) was injected at 0730 hours in the cephalic vein in awake animals on GD 43, 44 and 46 (Supplemental Table S3). Health and behavior observations were conducted three times pre-treatment, 6 hours after each of the three injections, and three times post-treatment. The checklist captured the presence or absence of any clinical or behavioral symptoms resulting from the infusions including change in appetite, watery eyes or nasal discharge, liquid stool, lethargy or labored movements, and body temperature. Prior to infusions, programmable temperature microchips (Bio Medic Temperature Systems, Seaford, DE) were implanted subcutaneously under sedation near the left and right clavicle, and a temperature wand then scanned the microchip and displayed body temperature. Temperatures were recorded just prior to Poly ICLC or saline injection, and 30 minutes, 6 hours, and 8 hours after injection. Pre- and post-treatment baseline temperatures were taken during health and behavior observations. Blood was collected from the dams on approximately GD 40 while sedated for ultrasound (pre-treatment), from awake animals on GD 44 and 46, six hours after Poly ICLC infusion, and on GD 51 or 52 while sedated for a recheck ultrasound (post-treatment) for cytokine analysis. Blood samples were centrifuged, and the serum was removed, aliquoted into 200 uL samples, and frozen at -80°C until analysis.

**Supplemental Table S3. Maternal Response to Poly ICLC**

| **Gestational Day (GD)** | **Poly ICLC Injection** | **Temperature** | **Appetite** | **Cytokine** |
| --- | --- | --- | --- | --- |
| Baseline (minimum 24 hours before first injection) | --- | Baseline  Temperature | 1:30pm  Three assessments  across 10 days | 8am  Baseline blood draw |
| GD 43 | 7:30am  Injection #1 | 7:30am  8:00am  1:30pm  3:30pm | 1:30pm | --- |
| GD 44 | 7:30am  Injection #2 | 7:30am  8:00am  1:30pm  3:30pm | 1:30pm | 1:30pm  Blood drawn 6 hours after injection #2 |
| GD 45 | No injection | --- | --- | --- |
| GD 46 | 7:30am  Injection #3 | 7:30am  8:00am  1:30pm  3:30pm | 1:30pm | 1:30pm  Blood drawn 6 hours after injection #3 |
| Baseline  (minimum 96 hours after last injection) | --- | Baseline Temperature | 1:30pm  Three assessments  across 10 days | 8am  Baseline blood  draw |

**Cytokine analysis.**

A longitudinal analysis on the maternal cytokine response to Poly ICLC exposure was measured in serum collected at baseline (pre-exposure) and after the second (GD 44) and third (GD 46) Poly ICLC injections. Using a nonhuman primate multiplexing bead immunoassay (Milipore-Sigma, Burlington, MA), we determined the levels IL-6. The kit was run according to the manufacturer’s instructions. Briefly, 25 µL of sample was incubated with antibody-coupled fluorescent beads and then washed and incubated with biotinylated detection antibodies followed by streptavidin–phycoerythrin. The beads were then analyzed using flow-based Luminex™ 100 suspension array system (Bio-Plex 200; Bio-Rad Laboratories, Inc.). Standard curves were generated by Bio-plex Manager software to determine unknown sample concentration, with reference cytokines provided by the manufacturer in the kit. The bead sets were analyzed using a flow-based Luminex™ 100 suspension array system (Bio-Plex 200; Bio-Rad Laboratories). A five-parameter model was used to calculate final concentrations. Concentrations obtained below the sensitivity limit of detection (LOD) were calculated as LOD/2 for statistical comparisons. Supernatant aliquots were free of any previous freeze/thaw cycles.

**Rearing conditions and husbandry.**

Infants were raised in individual cages with their mothers, where they had visual access to other mother-infant pairs at all times. For 3 hours each day, one familiar adult male and four familiar mother-infant pairs were allowed to freely interact in a large cage (3m l x 1.8m w x 2m h) to provide enrichment and facilitate species-typical social development. The rearing groups consisted of two MIA-treated mother-infant dyads and two control mother-infant dyads. Dominance hierarchies naturally formed between the dams, and group stability was monitored throughout by trained observers. The infants were weaned from their mothers at 6 months of age and were permanently paired with a familiar peer from their rearing group. Weanlings continued the same socialization routine through approximately 18 months of age. They were transferred to the large enclosures for 3 hours each day with the same three weanlings from their rearing group, the familiar adult male, and an adult female who was not one of the dams involved in the study. Animal rooms were maintained at 17.78–28.89°C and on a 12/12 light/dark cycle (lights on at 0600). Subjects were fed twice daily (Lab Diet #5047, PMI Nutrition International INC, Brentwood, MO), provided with forage scratch daily and fresh produce biweekly; they had access to water *ad libitum* and a variety of enrichment devices.

**Offspring physical growth.**

Measures of body growth were collected at 1 and 3 months of age as well as at neuroimaging time points (6, 12 and 24 months of age). Body measurements included weight, crown-rump length, and circumference of the head. Crown-rump length was measured as the distance from the top of the head to the base of the tail using a rigid board with embedded ruler. Head circumference was measured with a flexible tape measure around the widest portion of the skull.

**Offspring neurodevelopmental milestones and early behavioral development.**

The nonhuman primate offspring described in this paper are undergoing comprehensive assessments of social and cognitive development that will be the focus of future publications. Here we present data on a neurobehavioral neonatal assessment (1 week) and home cage observations of the mother-infant dyad (0-6 months), home cage observations with their age/sex/treatment matched cage mate (6-18 months) and introduction to a novel conspecific (11 months) to provide an assessment of species-typical developmental milestones. All behavioral observations were carried out by trained observers demonstrating an inter-observer reliability > 85% (agreements/ [agreements + disagreements] X 100). Each infant received a dye mark, allowing the observers to record behaviors while remaining blind to their experimental condition.

*Neonatal assessment* ***-*** The development of reflexes and basic neuromotor functioning was assessed at 7 (±1) days of age. The dam was lightly sedated with ketamine to remove the infant, and the infant was transferred to a testing room. A five point scale of maturity was used to rate the infant on measures of visual orientation and following; reflexes including rooting, righting, placing and Moro; motor maturity including head posture, coordination of movements and prone progression; and state control (agitation and consolability). For each assessment, infants were given a score of 2: present/fully developed; 1.5: mostly present/developed; 1: partially present or partially developed; 0.5: slightly developed or present; or 0: absent.

*Mother-infant home cage observation (0-6 months) -* Each mother-infant dyad was observed five days per week in the home cage by a trained observer who was familiar to the animals yet blinded to the condition of the animals. Interactions were quantified using a checklist of behaviors to describe the type and frequency of interactions between the mother and infant. Behaviors included nursing, grooming, contact, maternal behaviors (restrain, retrieve, rejection, aggression), facial expressions, environmental exploration, and stereotypy. Half of the dyads were observed in the morning and half were observed mid-day beginning when dyads were relocated to the project room and continued until weaning at approximately 6 months of age. The one-minute observation was broken into 6 10-second bins, and one-zero sampling was used to record behaviors. In one-zero sampling, every behavior that is present within each bin receives a score of 1 regardless of the number of times it occurs within the bin. Behaviors that are absent within each bin receive a score of 0.

*Infant-infant home cage observations (6-18 months) -* Each infant-infant dyad was observed three days per week for 52 weeks in the home cage by a trained observer who was familiar to the animals yet blinded to the condition of the animals. Interactions were quantified using a checklist of behaviors to describe the type and frequency of interactions between familiar peers. Behaviors included sleep, nonsocial activity, proximity, contact, play, environmental exploration, and stereotypy. Half of the dyads were observed in the morning and half were observed mid-day beginning one week after the dyads were formed at weaning at approximately 6 months of age. The one-minute observation was scored in 6 10-second bins as described above.

*Social Interest.* At approximately 11 months of age, social interactions with a novel conspecific were evaluated using a modified version of the rodent 3-chambered social approach assay. This assay provides a screen for sociability as indexed by the amount of time spent in the chamber containing the novel animal as well as time spent in proximity and contact with the novel animal. The focal animal is placed in a chute with visual access to two adjacent large chambers (5’w x 14’l x 7’h), connected by door, that each contain a holding cage. The holding cage in the “social chamber” contains a novel age- and sex-matched conspecific while the holding cage in the “nonsocial chamber” is empty. The focal animal is released from the chute by simultaneously opening two doors from the chute, one that accesses the social chamber and one that accesses the nonsocial chamber. The focal animal chooses which chamber to enter, the chute doors are closed, and the 15 minute session is video recorded for later scoring of time spent in the social chamber (containing the novel conspecific in a small holding cage) and the nonsocial chamber (containing an identical empty holding cage). While in the social chamber, the durations of time spent in the immediate proximity (i.e., within arms reach) or contact with the stimulus animal was recorded, as was the amount of time spent away (i.e., more than arm’s reach distance) from the stimulus. The frequency of establishing proximity and the number of entries into the social chamber were also quantified.

**Supplemental Table S4.** Home Cage Interaction Behavioral Ethogram

| ***Behavior*** | ***Description*** |
| --- | --- |
| *Maternal Contact (pre-wean only)* | |
| Breast Contact | Focal infant suckles mother. |
| Ventral Contact | Ventral surface of the focal contacts ventral surface of dam. |
| Other Contact | Other physical contact (not breast contact or ventral contact) with dam. |
| No Contact | No physical contact between infant and dam. |
| *Maternal Interaction (pre-wean only)* | |
| Maternal Restrain | Mother physically interferes with the infant’s attempts to move away from her. |
| Maternal Retrieve | Mother physically brings infant closer to her. |
| Maternal Reject | Mother physically prevents the focal infant from contact. |
| *Exploratory Events (composite score)* | |
| Toy Play | Oral or manual manipulation of toys in cage. |
| Oral Explore | Oral manipulation to any part of the cage excluding food. |
| Manual Explore | Manual manipulation to any part of the cage. |
| *Total Stereotypies (composite score)* | |
| Pace | Repetitive undirected pacing with the same path repeated at least 3 consecutive times. |
| Head Twist | Throwing of the head back and to the side in an exaggerated manner. |
| Backflip | Repetitive backflip at least two times in a row. |
| Bounce | Repetitive bounce up and down at least two times in a row. |
| Nipple Clasp | Holding of the nipple. |
| Rock | Stationary rocking either back and forth or side to side. |
| Swing | Swinging within the cage for at least 3 sec. |
| Self-bite | Biting motion of own limb or body part. |
| Salute | Fingers or hand held in place along the brow, eyes or other part of the upper face. |
| Other Abnormal Behavior | Any other abnormal behaviors not described above. |
| *Infant-Infant Interaction (post-wean only)* | |
| Non-Social Activity | Not in proximity, contact, play or other social activity. |
| Home cage proximity | Both animals are in the same cage. |
| Contact | Any physical contact between the focal animal and another. |
| Play | Any instance of play (contact play, wrestle play, chase). |

**Neuroimaging**.

Magnetic resonance imaging was performed at approximately 6, 12 and 24 months of age using a Siemens Magnetom Skyra 3-T (Davis, California) with 8-channel coil optimized for monkey brain scanning (RapidMR, Columbus, Ohio). Twenty-four of the animals were also scanned at 1 month of age. However, due to the low gray matter/white matter contrast in the T1 images, we elected not to include this time point in the analyses for the current paper. Three animals were scanned at 3 months of age. However, we encountered some respiratory difficulties at this age and we elected to stop scanning in order to not put the animals at risk. We determined that three of the animals were sensitive to isoflurane. At subsequent time points, these animals were scanned using propofol as the anesthetic. The rate of infusion varied in order to maintain the animal at a steady state of anesthesia. All other animals at all time points were sedated with ketamine for tracheal intubation then anesthetized with Isoflurane for positioning in an MR-compatible stereotaxic apparatus. Once the animal was placed in and centered at the mid-line of the stereotaxic apparatus, the 8-channel receiver coil was attached to the stereotaxic apparatus using a custom connector. The center point of the 8-channel coil was positioned at AP+10 on the stereotaxic apparatus. The Skyra table was “landmarked” at AP+10 so that the center of the animal’s brain was at the isocenter of the MRI magnet. Anesthesia was maintained with isoflurane at 1.3-2.0%. Fluids were maintained with a saline infusion at a rate of 10 ml/kg/hr for the duration of the MRI scan.

*Acquisition Parameters*: T1 weighted images (480 sagittal slices) were acquired with TR=2500 ms, TE=3.65 ms, flip angle=7°, field of view 256x256, voxel size during acquisition 0.6x0.6x0.6 mm. Acquired images were interpolated during image reconstruction to 512x512 voxels with a final resolution of 0.3 x 0.3 x 0.3 mm. The structural imaging protocol was followed with additional sequences that will not be described in this paper.

*Image processing*: All images were processed blinded to group assignment. T1 weighted images were aligned into common ^2^, bias field corrected, brain masked using AutoSeg_3.3.2 ^3^. Brain masks were manually corrected if necessary. Following this preprocessing T1 weighted images were segmented into gray matter (GM), white matter (WM) and cerebrospinal fluid (CSF) using NeosegPipeline_v1.0.8 ^4^(Supplemental Figure S1). Probabilistic tissue maps from structural multi-atlas templates were applied to each subject’s T1 weighted images via deformable registration University of North Carolina (UNC) lobar parcellation were employed to parcellate the tissue segmentations into 24 lobar brain regions using the multi-atlas fusion in AutoSeg_3.3.2 ^3^ (See Supplemental Figure S1). For these regions, total GM and WM volumes were extracted. Lateral ventricle volumes were determined via semi-automated segmentation using the region competition deformable surface approach in ITK snap ^5^ as applied to the probability CSF maps from the tissue segmentation ^6^. All segmentation and parcellation results were visually quality controlled. No major issues were detected for any of the results.

*Regions of interest*. In order to limit the number of comparisons , we selected 5 regions of interest based on a review of literature documenting brain alterations associated with MIA: prefrontal, frontal, cingulate, temporal limbic (including amygdala and hippocampus) regions (Supplemental Figure S2), and lateral ventricles ^7-11^.

**Supplemental Figure S1**. (A) Structural MRI analysis workflow. GM - gray matter, WM - white matter. (B) Multi-atlas parcellation gives more accurate results than single-atlas parcellation: rows show regions that were improved with multi-atlas parcellation (insula and temporal limbic).

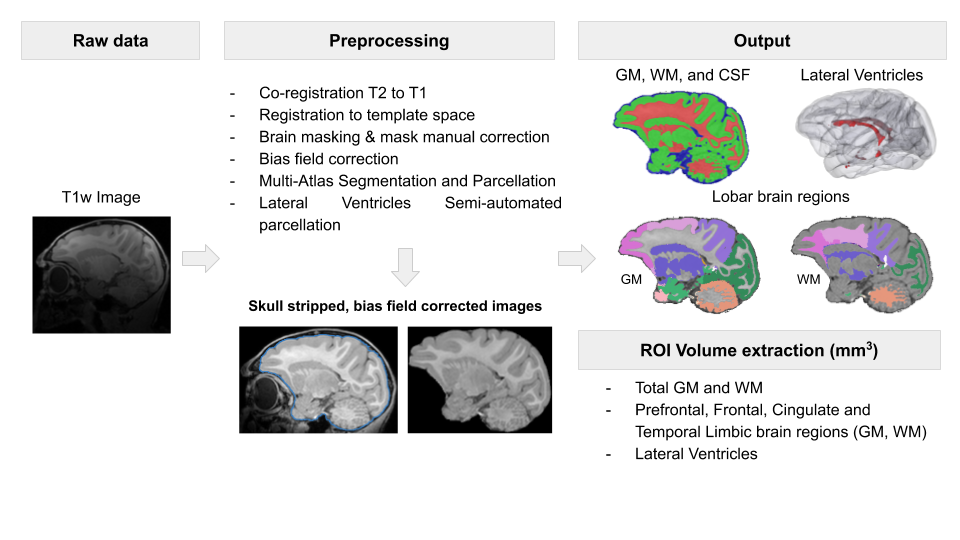

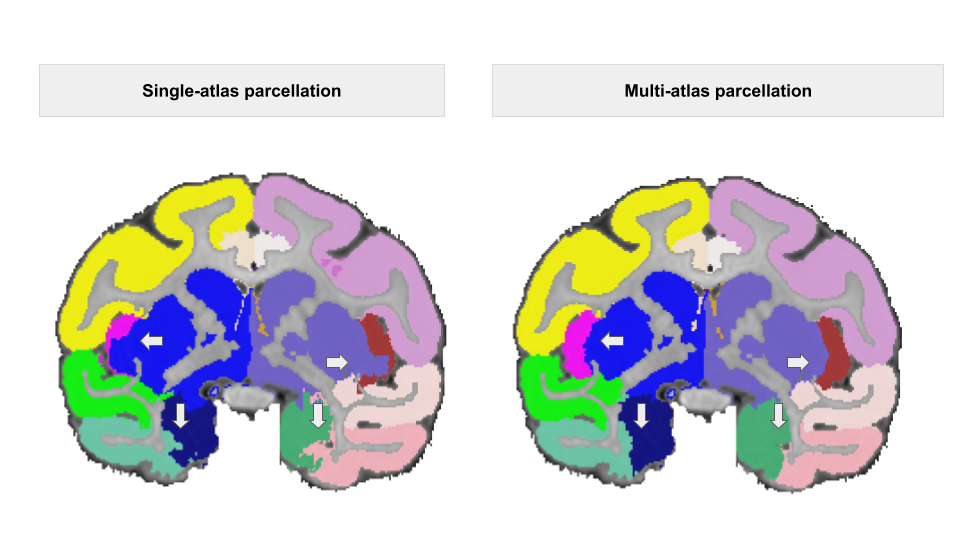

Supplemental **Figure S2**. Regions of interest visualization

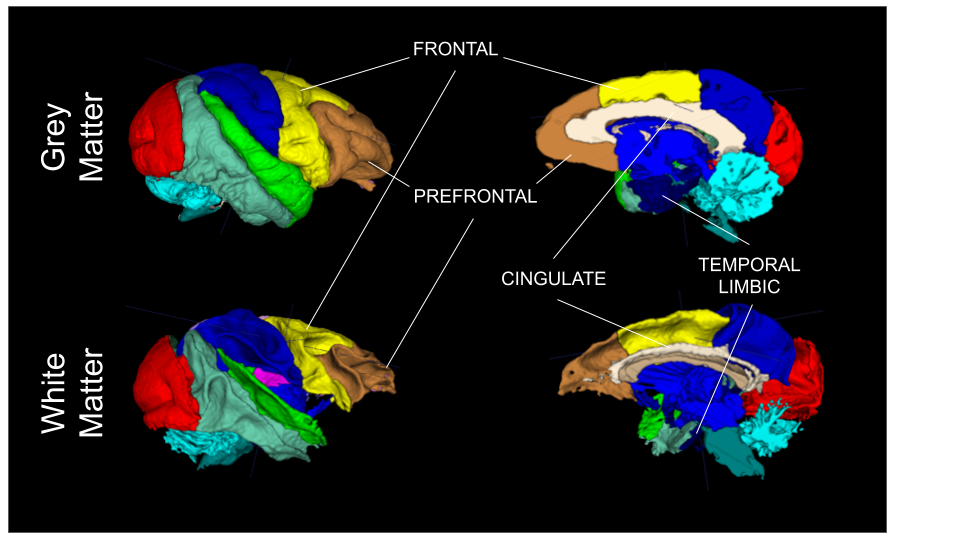

**Statistical analysis.**

*Offspring development analyses.* Developmental trajectories for weight, crown-rump and head circumference were analyzed using mixed-effects models with fixed effects for group (MIA, control), age at measurement, and the interaction between age and group. Linear, quadratic, and cubic age effects were considered. To account for the within-animal dependence, random effects for animal, linear and quadratic effect of age were included in the models.

*Social Interest.* The age at which infants were observed ranged from 9.4 to 11 months and there was a statistically significant difference in age at testing between MIA and control group (*p* = .048). Preliminary analyses revealed significant effects of age at testing, and for some behaviors the effect of age in MIA differed from the one in control infants. Thus, for each behavior duration or frequency, we fitted a general linear model that included fixed effects for group (MIA, control), stimulus animal, age at testing (centered at the median, 10.2 months), and the interaction of group with age. The estimate for the group effect in these models can be interpreted as the estimated difference between 10.2-month old MIA and Control animals tested with the same stimulus animal. All models had heterogeneous variances to account for the higher variability in the control group. Some of the variables were square-root transformed to improve normality.

*MRI analyses.* The goals of the statistical models were to estimate patterns of change in the outcome variables from 6 to 24 months and to test whether MIA-treated animals had a different developmental trajectory than the control animals. Separate models were fitted for each of the global measures (brain volume, GM, WM, left and right ventricles). We first fitted models with fixed effects for group (MIA, control), time (6, 12 or 24 months) and interaction between group and time, as well as a random effect for animal, to account for correlated nature of the data due to the repeated measures. If the interaction did not add significantly to the model, then it was removed and the results of the model including only main effects were reported. WM and GM ROI volume data were doubly multivariate, with measurements taken at 3 time points on each side of the brain (left, right). To further account for the correlations between left and right side measurements, we used separable structure (UN@CS) to model these data. In these analyses, separate models were fitted for WM and GM volumes in frontal, prefrontal, cingulate, and temporal limbic cortices. The core models included fixed effects for group (MIA, control), time (6, 12 or 24 months), side (left, right), and interaction between group and time, and a random effect for animal. We removed the interaction term from the reported models if it was not significant. A second set of models was fitted by adding a term for the total brain.

**SUPPLEMENTAL RESULTS**

**Validation of maternal immune activation**

Appetite (Supplemental Table S5), temperature (Supplemental Table S6; Supplemental Figure S3) and cytokine profiles (Table 1; Supplemental Figure S4) of the pregnant monkeys confirmed a strong inflammatory response to Poly ICLC.

**Table S5**. Descriptive statistics for appetite changes during Poly ICLC injections.

|  | **MIA**  **(*n* = 14)** | | **Control**  **(*n* = 10)** | |
| --- | --- | --- | --- | --- |
| ***Time*** | **Mean (*SD*)** | **Median [Range]** | **Mean (*SD*)** | **Median [Range]** |
| Pre-treatment^a^ | 1.1 (0.7) | 1.3 [0 – 2] | 1.5 (0.4) | 1.5 [0.7 – 2] |
| Treatment | 0.4 (0.4) | 0.3 [0 – 1.3] | 1.6 (0.4) | 1.7 [0.7 – 2] |
| Post-treatment^b^ | 1.3 (0.5) | 1.3 [0.3 – 2] | 1.3 (0.5) | 1.3 [0.3 – 2] |

Abbreviations: MIA, maternal immune activation; SD, standard deviation; GD, gestation day.

Pre-treatment = 3 days between GD 32 and 42; Treatment = GD 43, 44, and 46; Post-treatment = 3 days between GD 47 and 57.

Appetite was rated on a 0-2 Likert-type scale, with: 0 = *Poor (1-3 biscuits eaten)*, 1 = *Fair (4-6 biscuits eaten)*, 2 = *Good (7-9 biscuits eaten)*. For each period, scores were first summarized within-animal, by calculating the average over three days.

^a^All three days missing data for 1 animal in MIA group; ^b^One day missing data for 1 animal in MIA group

**Supplemental Table S6.** Descriptive statistics (mean, standard deviation) for temperature (^o^F) changes during Poly ICLC (MIA) or saline (CON) injections.

|  | **Gestation Day 43** | | **Gestation Day 44** | | | | **Gestation Day 46** | |
| --- | --- | --- | --- | --- | --- | --- | --- | --- |
|  | **MIA**  **(*n* = 14)** | **Control**  **(*n* = 10)** | | **MIA**  **(*n* = 14)** | **Control**  **(*n* = 10)** | **MIA**  **(*n* = 14)** | | **Control**  **(*n* = 10)** |
| Time-point for temperature reading | | | | | | | | |
| Pre-infusion | 97.9 (1.2) | 97.2 (2.9) | | 98.1 (1.5) | 98.1 (1.3) | 97.9 (1.1) | | 98.4 (1.1) |
| 30 minutes | 98.2 (1.6) | 98.5 (1.3) | | 97.6 (1.5) | 98.4 (1.0) | 97.9 (1.0) | | 98.8 (1.1) |
| 6 hours | 100.2 (1.1) | 97.7 (1.3) | | 99.7 (1.5) | 97.9 (0.9) | 100.1 (1.5) | | 98.5 (1.2) |
| 8 hours | 99.2 (1.0) | 97.8 (1.4) | | 99.2 (1.8) | 97.8 (1.2) | 98.6 (2.1) | | 97.7 (1.9) |

Abbreviations: MIA, maternal immune activation; SD, standard deviation

^a^Pre-infusion temperatures were recorded immediately prior to the injection (within 1-2 minutes).

**Supplemental Figure S3**. Average temperature for the MIA and saline treated dams from pre-infusion (0) through 480 minutes after infusion during gestation days 43, 44 and 46. Vertical bars represent 1 standard deviation.

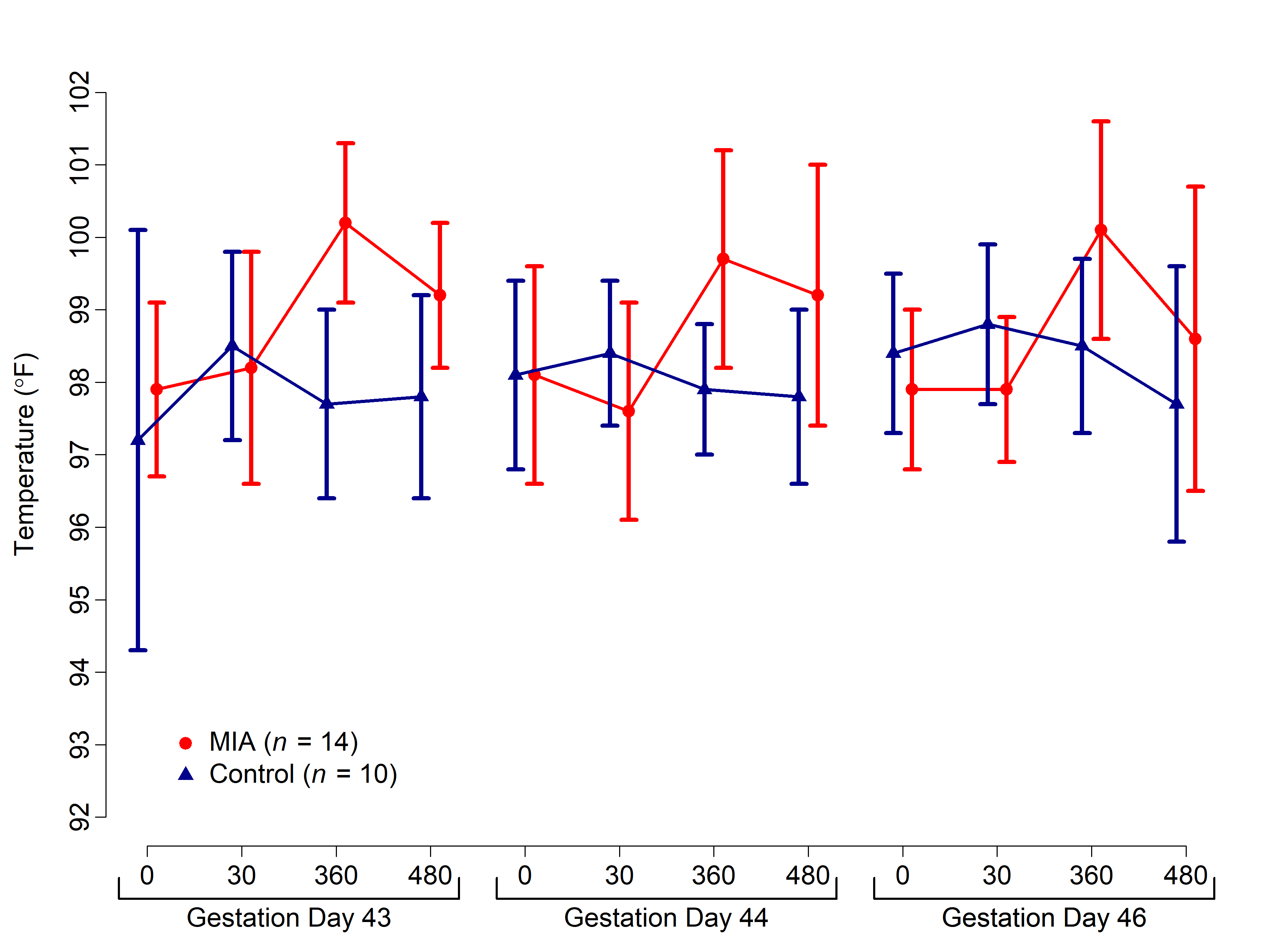

**Supplemental Figure S4**. IL-6 response for the MIA (a) and saline (b) treated dams. Baseline cytokine levels (pre-dosing GD 40), 6 hours after the second Poly ICLC injection (GD 44) and 6 hours after the third PolyIC injection (GD 46). To aid visualization, data were square root transformed. Poly ICLC was associated with a transient increase in IL-6 after the 2nd and 3rd injections. Black arrows represent the 2nd and 3rd Poly ICLC injections time points (blood samples were not collected after the first Poly ICLC injection to minimize stress to the dam).

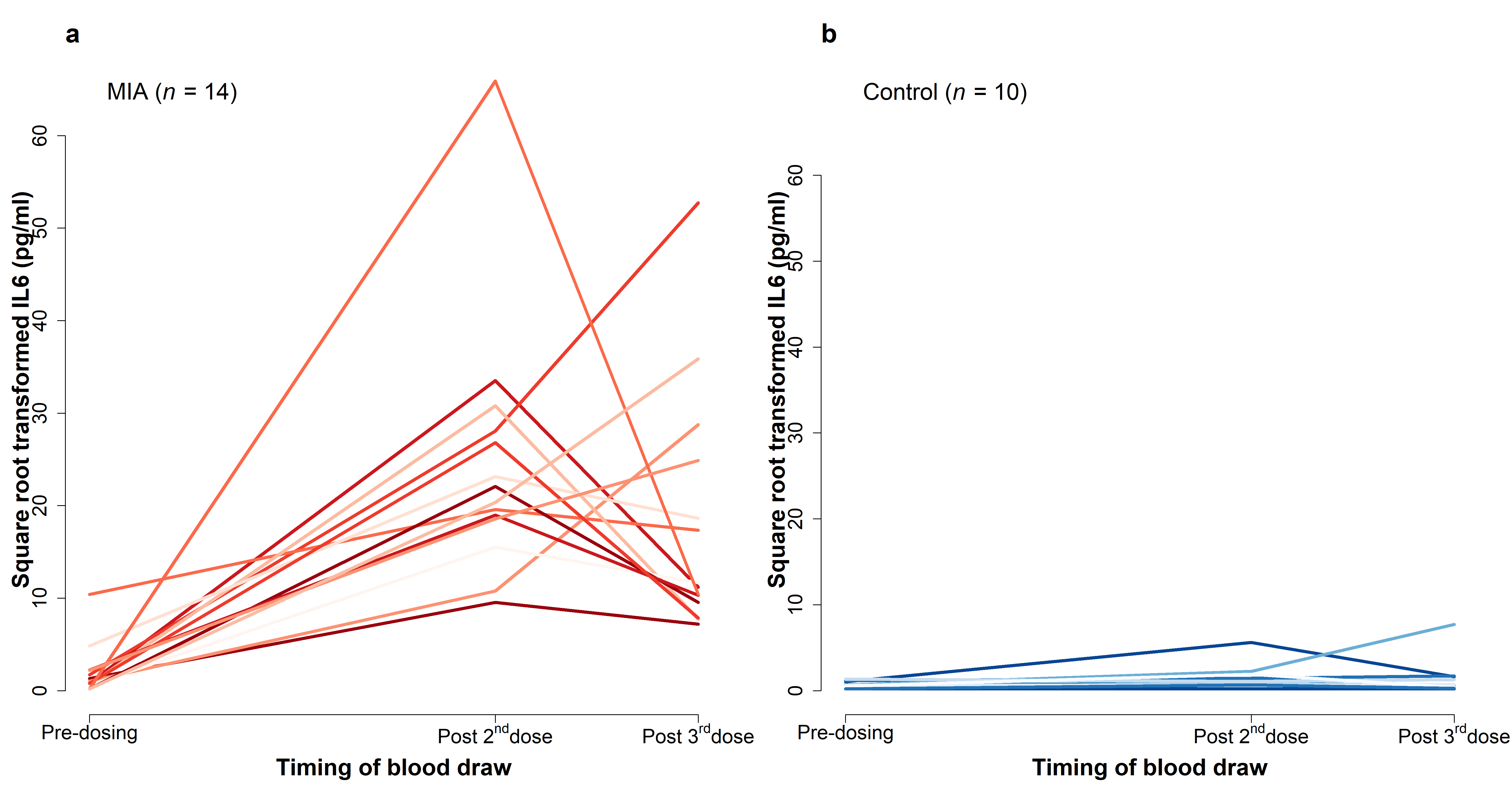

**Offspring development**

There were no group differences in overall health, physical development (Supplemental Table S7) and neuromotor-reflexes, behavioral maturation and attention (Supplemental Table S8). There were also no significant group differences detected in the home cage observations with their mothers from 9-24 weeks of age (Supplemental Table S9) or when observed interacting with a treatment-matched social partner in the home cage from 34-78 weeks of age (Supplemental Table S10).

**Supplemental Table S7**. Summary for the morphometric measures from 1 to 24 months

|  | Age (days)  *mean* (*SD*) [*Range*] | | Weight (kg)  *mean* (*SD*) [*Range*] | | | Crown Rump (cm)  *mean* (*SD*) [*Range*] | | | | Head Circumference (cm)  *mean* (*SD*) [*Range*] | | | |
| --- | --- | --- | --- | --- | --- | --- | --- | --- | --- | --- | --- | --- | --- |
|  | MIA | Control | MIA | | Control | | MIA | Control | | | MIA | Control | |
| Evaluation time (months) | | | | | | | | | |  | | | |
| 1^a^ | 31 (2)  [28-35] | 32 (7)  [28-54] | 0.7 (0.1)  [0.5-0.8] | 0.7 (0.2)  [0.4-1.0] | | 22.4 (1.2)  [20.5-24.5] | | | 21.9 (1.3)  [19.3-24.0] | 20.8 (0.7)  [19.5-21.8] | | | 21.1 (0.8)  [19.5-22.4] |
| 3 | 91 (2)  [88-94] | 91 (1)  [88-92] | 1.0 (0.1)  [0.8-1.3] | 1.0 (0.3)  [0.6-1.7] | | 25.1 (1.4)  [22.7-28.0] | | | 24.7 (1.9)  [20.2-28.5] | 22.2 (0.9)  [20.9-23.9] | | | 22.5 (1.0)  [20.2-24.1] |
| 6 | 180 (2)  [177-182] | 180 (1)  [177-181] | 1.5 (0.2)  [1.3-1.8] | 1.5 (0.2)  [1.1-1.8] | | 30.1 (1.8)  [27.0-33.0] | | | 30.2 (2.1)  [27.0-33.5] | 23.4 (0.6)  [22.5-24.5] | | | 23.9 (1.2)  [21.2-26.0] |
| 12^b^ | 365 (2)  [364-369] | 365 (1)  [364-366] | 2.3 (0.3)  [1.9-2.9] | 2.3 (0.3)  [1.8-2.9] | | 35.8 (1.4)  [34.0-38.0] | | | 35.2 (3.1)  [30.0-39.0] | 24.7 (0.6)  [23.8-26.0] | | | 24.9 (0.8)  [23.5-26.5] |
| 24^b^ | 730 (1)  [729-733] | 730 (1)  [729-732] | 3.7 (0.5)  [2.9-4.6] | 3.8 (0.6)  [2.8-4.7] | | 41.9 (2.0)  [38.1-45.0] | | | 41.9 (1.9)  [38.3-44.7] | 26.5 (1.0)  [24.5-28.6] | | | 27.2 (1.3)  [24.5-29.5] |

Abbreviations: MIA, maternal immune activation; SD, standard deviation.

^a^Data missing for 1 MIA and 2 Control animals for crown rump and 1 MIA and 1 Control for head circumference; ^b^Data missing for 1 MIA animal on all variables due to death.

**Supplemental Table S8.** Summary of week 1 neuro-motor reflexes, behavioral maturation and attention

|  | **MIA**  **(*n* = 14)** | | **Control**  **(*n* = 14)** | | ***P*-value^a^** |
| --- | --- | --- | --- | --- | --- |
|  | **Mean (*SD*)** | **Median [Range]** | **Mean (*SD*)** | **Median [Range]** |  |
| Age (days) | 7.0 (0.7) | 7.0 [6.0 – 8.0] | 6.9 (0.6) | 7.0 [6.0 – 8.0] | 0.95 |
| Weight (kg)^b^ | 0.5 (0.1) | 0.5 [0.5 – 0.7] | 0.6 (0.1) | 0.6 [0.4 – 0.7] | 0.49 |
| Gestation length (days) | 168 (4) | 167 [162 – 175] | 167 (4) | 167 [160 – 177] | 0.97 |
| Visual Orientation | 1.3 (0.6) | 1.3 [0 – 2] | 1.7 (0.4) | 1.8 [1 – 2] | 0.14 |
| Visual Follow | 0.9 (0.4) | 1.0 [0.5 – 1.5] | 1.1 (0.6) | 1.0 [0 – 2] | 0.23 |
| Head Posture | 2.0 (0.0) | 2.0 [2 – 2] | 2.0 (0.0) | 2.0 [2 – 2] | 1.00 |
| Coordination | 0.8 (0.8) | 0.8 [0 – 2] | 0.7 (0.7) | 0.8 [0 – 2] | 0.96 |
| Spontaneous Crawl | 1.1 (0.7) | 1.0 [0 – 2] | 0.9 (0.7) | 1.0 [0 – 2] | 0.38 |
| Rooting | 0.6 (0.9) | 0.0 [0 – 2] | 0.4 (0.7) | 0.0 [0 – 2] | 0.78 |
| Righting | 1.9 (0.5) | 2.0 [0 – 2] | 1.9 (0.5) | 2.0 [0 – 2] | 1.00 |
| Placing | 1.1 (0.8) | 1.0 [0 – 2] | 0.8 (0.8) | 0.8 [0 – 2] | 0.36 |
| Moro Reflex^c^ | 1.8 (0.3) | 2.0 [1 – 2] | 1.7 (0.4) | 2.0 [1 – 2] | 0.76 |
| Predominant State | 0.9 (0.5) | 1.0 [0 – 2] | 0.6 (0.6) | 0.5 [0 – 2] | 0.19 |
| Consolability | 0.5 (0.7) | 0.0 [0 – 2] | 0.3 (0.6) | 0.0 [0 – 1.5] | 0.36 |

Abbreviations: MIA, maternal immune activation; SD, standard deviation.

Note: All animals were rated with a score from 0 to 2 (0 = reflex absent, 0.5 = reflex slightly developed or present, 1 = partially present or partially developed, 1.5 = mostly present or developed, 2 = reflex present or fully developed). Predominant State: 0 = calm, alert, aware, 0.5 = mostly calm with slight agitation, 1 = alert but agitated for no more than half the exam, 1.5 = agitated for more than half the exam, 2 = extremely agitated throughout entire exam. Consolability: 0 = quickly consoled when picked up following exam, 0.5 = consoled after brief period of holding and swaddling, 1 = infant consoled only after prolonged holding, swaddling, rocking and/or stroking, 1.5 = brief moments of consolation and quiet after prolonged holding, 2 = inconsolable.

^a^From Wilcoxon two-sample exact tests; ^b^Data missing data for 2 animals in MIA group; ^c^Data missing for 1 animal in MIA group.

**Supplemental Table S9.** Summary of pre-wean (weeks 9 to 24) home cage observations

|  | **MIA**  **(*n* = 14)** | | **Control**  **(*n* = 14)** | | ***P*-value** |
| --- | --- | --- | --- | --- | --- |
|  | **Mean (*SD*)** | **Median [Range]** | **Mean (*SD*)** | **Median [Range]** |  |
| Breast contact | 2.4 (0.8) | 2.6 [1.2 – 3.6] | 2.7 (0.8) | 2.7 [1.5 – 3.9] | 0.36 |
| Other contact | 0.7 (0.3) | 0.6 [0.2 – 1.2] | 0.8 (0.5) | 0.7 [0.2 – 2.1] | 0.96 |
| Ventral contact | 1.5 (0.9) | 1.3 [0.4 – 3.8] | 1.5 (0.6) | 1.4 [0.5 – 2.6] | 0.98 |
| No contact | 3.4 (0.9) | 3.4 [1.8 – 4.8] | 3.2 (0.8) | 3.1 [2.0 – 4.4] | 0.48 |
| Maternal restrain | 0.02 (0.04) | 0.1 [0 – 0.13] | 0.01 (0.02) | 0 [0 – 0.07] | 0.08 |
| Maternal retrieve | 0.04 (0.05) | 0.02 [0 – 0.19] | 0.03 (0.03) | 0.03 [0 – 0.10] | 0.88 |
| Maternal reject | 0.11 (0.12) | 0.07 [0 – 0.44] | 0.08 (0.08) | 0.06 [0 – 0.31] | 0.68 |
| Total explore^a^ | 0.4 (0.2) | 0.4 [0.1 – 0.7] | 0.4 (0.1) | 0.4 [0.2 – 0.6] | 1.00 |
| Total stereotypies^b^ | 0.003 (0.01) | 0 [0 – 0.3] | 0.004 (0.1) | 0.0 [0 – 0.2] | 0.52 |

Abbreviations: MIA, maternal immune activation; SD, standard deviation.

^a^Total explore includes toy play, oral explore, manual explore; ^b^Total stereotypy includes pace, head twist, backflip, bounce, nipple clasp, rock, swing, self-bite, salute.

Note: Infants were observed up to 5 times per week between 2 and 6 months. For each animal and each week, behaviors were first averaged over the observations that were available (ranging from 3 to 5). All animals had data between 9 and 24 weeks, so we then averaged the behaviors again within animals (from 9 to 24 weeks), to create a summary over the course of the study. Wilcoxon two-sample exact tests were then fitted to these averaged behaviors.

**Supplemental Table S10.** Summary of post-wean (weeks 34 to 78) home cage observations

|  | **MIA**  **(*n* = 14)** | | **Control**  **(*n* = 14)** | | **MIA vs. Control Estimated Difference (*SE*)** | ***P*-value**^a^ |
| --- | --- | --- | --- | --- | --- | --- |
|  | **Mean (*SD*)** | **Median [Range]** | **Mean (*SD*)** | **Median [Range]** |  |  |
| Sleep^b^ | 0.03 (0.05) | 0 [0 – 0.1] | 0.04 (0.03) | 0.04 [0 – 0.1] | -0.07 (0.05) | 0.20 |
| Non-social^b^ | 3.3 (0.6) | 3.5 [2.2 – 4.3] | 3.4 (0.6) | 3.4 [2.0 – 4.1] | 0.04 (0.04) | 0.30 |
| Proximity | 4.0 (0.5) | 4.0 [3.3 – 4.7] | 3.9 (0.5) | 3.8 [3.1 – 4.8] | 0.07 (0.17) | 0.68 |
| Contact^b^ | 0.5 (0.2) | 0.5 [0.1 – 0.8] | 0.5 (0.2) | 0.5 [0.1 – 1.1] | 0.01 (0.06) | 0.92 |
| Play^b^ | 1.0 (0.3) | 0.9 [0.5 – 1.4] | 0.9 (0.3) | 0.8 [0.5 – 1.4] | -0.05 (0.04) | 0.31 |
| Composite activity^b^ | 0.5 (0.3) | 0.4 [0.2 – 1.1] | 0.6 (0.3) | 0.5 [0.2 – 1.1] | -0.04 (0.08) | 0.67 |
| Composite stereotypies^b^ | 0.2 (0.3) | 0.1 [0 – 0.8] | 0.2 (0.3) | 0.1 [0 – 1.0] | 0.02 (0.12) | 0.87 |

Abbreviations: MIA, maternal immune activation; SD, standard deviation; SE, standard error.

^a^From linear mixed effect models; ^b^Data was square root transformed for the analysis.

Note: Infants were observed 3 times per week between 27 and 85 weeks. For each animal and each week, behaviors were first averaged over the 3 observations that were available. Not all animals were observed each week; all animals had data between 34 and 78 weeks, so we then averaged each type of behavior again within animals (from 34 to 78 weeks), to create summary behaviors over the course of the study. To these summary behaviors we fitted linear mixed-effects models with a fixed effect for group and a random effect for the “play buddy” to account for the fact that animals interacted in pairs and the exhibited behaviors were correlated.

**Neuroimaging.**

**Supplemental Table S11.** Parameter estimates from the linear mixed-effects models for global volumetric measures

|  | **Brain** | | **Gray** | | | | **White** | | **Left Ventricle** | | | **Right Ventricle** | | |
| --- | --- | --- | --- | --- | --- | --- | --- | --- | --- | --- | --- | --- | --- | --- |
| **Model term** | **Estimate**  **(*SE*)** | ***P*** | | **Estimate**  **(*SE*)** | ***P*** | **Estimate**  **(*SE*)** | | ***P*** | | **Estimate (*SE*)** | ***P*** | | **Estimate (*SE*)** | ***P*** |
| Intercept | 86536 (2393) | <.001 | | 63413 (1633) | <.001 | 23123 (801) | | <.001 | | 247 (21) | <.001 | | 259 (18) | <.001 |
| Difference (mm^3^) *MIA* vs. *Control* | -5504 (3432) | .12 | | -3862 (2339) | .11 | -1642 (1146) | | .16 | | -20 (29) | .50 | | -12 (25) | .63 |
| Difference (mm^3^) Time 2 vs. Time 1 | 2263 (418) | <.001 | | -247 (311) | .43 | 2610 (166) | | <.001 | | -6 (10) | .56 | | -7 (11) | .52 |
| Difference (mm^3^) Time 3 vs. Time 1 | 7279 (418) | <.001 | | 1411 (311) | <.001 | 5867 (166) | | <.001 | | 26 (10) | .01 | | 18 (11) | .09 |

Note: MIA = maternal immune activation, SE = standard error, Time 1 = 6 months, Time 2 = 12 months, Time 3 = 24 months. Mixed-effects linear regression models were fitted to 13 MIA and 14 control animals and included fixed effects for group and time and a random effect for animal. One additional animal was excluded from the Ventricle models due to extreme data. Interactions between group and time were added to the models but were not retained in the reported models because the overall tests for time by group were not significant. The intercept can be interpreted as the predicted Time 1 volume (in mm^3^) for a *Control* animal.

**Supplemental Table S12.** Parameter estimates from the unadjusted linear mixed-effects models for gray and white ROI volumetric measures

|  | **Gray Matter Volume** | | | | | | | |
| --- | --- | --- | --- | --- | --- | --- | --- | --- |
| **Model term** | **Frontal** | | **Prefrontal** | | **Cingulate** | | **Temporal Limbic** | |
|  | **Estimate (*SE*)** | ***P*** | **Estimate (*SE*)** | ***P*** | **Estimate (*SE*)** | ***P*** | **Estimate (*SE*)** | ***P*** |
| Intercept | 3719.7 (95.5) | <.001 | 3561.4 (112.0) | <.001 | 1181.3 (35.7) | <.001 | 1271.7 (32.8) | <.001 |
| Difference (mm^3^) *MIA* vs. *Control* | **-382.6 (135.8)** | **.009** | **-414.5 (160.4)** | **.02** | -95.0 (51.1) | .07 | -35.5 (45.9) | .45 |
| Difference (mm^3^) Time 2 vs. Time 1 | 68.7 (27.8) | .02 | -21.4 (22.5) | .34 | 1.1 (7.0) | .87 | 125.5 (11.1) | <.001 |
| Difference (mm^3^) Time 3 vs. Time 1 | 273.6 (27.8) | <.001 | 6.7 (22.5) | .77 | 1.1 (7.0) | .87 | 268.3 (11.1) | <.001 |
| Difference (mm^3^) Left vs. Right | 24.0 (13.4) | .09 | 11.9 (7.4) | .12 | -34.5 (9.2) | <.001 | -75.3 (7.3) | <.001 |
|  | **White Matter Volume** | | | | | | | |
| **Model term** | **Frontal** | | **Prefrontal** | | **Cingulate** | | **Temporal Limbic** | |
|  | **Estimate (*SE*)** | ***P*** | **Estimate (*SE*)** | ***P*** | **Estimate (*SE*)** | ***P*** | **Estimate (*SE*)** | ***P*** |
| Intercept | 1370.8 (48.6) | <.001 | 855.9 (41.4) | <.001 | 181.5 (8.4) | <.001 | 223.6 (7.6) | <.001 |
| Difference (mm^3^) *MIA* vs. *Control* at Time 1 | -92.7 (69.8) | .20 | -66.9 (59.7) | .27 | -22.5 (11.9) | .07 | -10.6 (10.7) | .33 |
| Difference (mm^3^) Time 2 vs. Time 1 in *Control* | 190.0 (14.6) | <.001 | 158.1 (12.9) | <.001 | 16.2 (2.1) | <.001 | 18.6 (2.1) | <.001 |
| Difference (mm^3^) Time 3 vs. Time 1 in *Control* | 454.7 (14.6) | <.001 | 353.2 (12.9) | <.001 | 46.7 (2.1) | <.001 | 56.7 (2.1) | <.001 |
| Difference between groups in Time 2 vs. Time 1 differences | -33.1 (21.0) | .12 | -26.5 (18.6) | .16 | – | – | – | – |
| Difference between groups in Time 3 vs. Time 1 differences | -70.9 (21.9) | .002 | -61.2 (18.6) | .002 | – | – | – | – |
| Difference (mm^3^) Left vs. Right | 31.8 (7.9) | <.001 | -10.7 (3.9) | .01 | 10.4 (2.2) | <.001 | -1.6 (2.1) | .46 |

Note: MIA = maternal immune activation, SE = standard error, Time 1 = 6 months, Time 2 = 12 months, Time 3 = 24 months. Mixed-effects linear regression models were fitted to 13 MIA and 14 control animals and included fixed effects for group, time and their interaction, side, and a random effect for animal. Interactions were not retained in the reported models if the overall test for time by group was non-significant. The intercept can be interpreted as the predicted Time 1 right volume (in mm^3^) for a *Control* animal.

Statistically significant MIA vs. Control group differences (*p* < .05) bolded for emphasis.
